# Revisiting the window of opportunity for co-transcriptional splicing efficiency and fidelity

**DOI:** 10.1101/497040

**Authors:** Vahid Aslanzadeh, Jean D. Beggs

## Abstract

Recently, we reported that changes in transcription elongation rate affect the efficiency and fidelity of precursor mRNA (pre-mRNA) splicing, especially of ribosomal protein (RP) transcripts. Here, we analyse these results in more detail, finding that the majority of RP transcripts with non-consensus 5’ splice sites have reduced splicing efficiency with faster transcription elongation, and improved efficiency with slower elongation, as might be predicted by the “window of opportunity” model for co-transcriptional splicing. In contrast, both faster and slower elongation reduce splicing fidelity, often for the same splicing events, and both faster and slower transcription increase fidelity with a different set of splicing events. We propose that certain non-consensus 5’ splice sites in ribosomal protein transcripts confer a stronger effect of transcription elongation rate on splicing efficiency, possibly by causing a rate-limiting step that delays activation of spliceosomes. The effects of different rates of transcription elongation on splicing fidelity are more difficult to explain by a simple window of opportunity model. We discuss these new findings in the context of current models of co-transcriptional splicing and splicing fidelity.

## Introduction

### Co-transcriptional splicing

The coding sequences of most eukaryotic genes are disrupted by introns (mainly non-coding sequences) that are also present in the nascent transcripts. Pre-mRNA splicing is the process that removes the introns and joins the coding segments (exons) to produce mature mRNAs. Evidence accumulating during the last two decades suggests that many, and possibly the majority of, splicing events occur co-transcriptionally, that is before transcription termination [1–7]. This raises the intruiging possibility of functionally significant interactions between splicing, chromatin, transcription and other RNA processing events, if they occur in close proximity [8–12]. Two models were proposed to explain how transcription and splicing are coupled; referred to as the recruitment coupling and the kinetic coupling models, which are not mutually exclusive (reviewed [13]).

### Recruitment coupling

The term “recruitment coupling” refers to the ability of the transcription machinery to promote the recruitment of RNA processing factors to the site of transcription. In particular, the carboxyterminal domain (CTD) of the largest subunit of RNA polymerase II (RNAPII) acts as a ‘landing pad’ for co-transcriptional recruitment of capping, splicing and 3’ end processing factors to nascent RNA [14]. Strong support for recruitment of splicing factors was obtained by a study with human cell lines showing CTD-dependent inhibitory action of serine/arginine-rich (SR) splicing factor SRSF3 (SRp20) in inclusion of fibronectin cassette exon 33 (E33)[15].

### Kinetic coupling

Early evidence for coupling between transcription and splicing was reported by Eperon et al. [16], who showed that, in HeLa cells, the use of an alternative splice site positioned within a potential stem-loop structure depended on a “window” of availability for splicing factors or hnRNP proteins to bind the nascent RNA before the stem-loop formed. A prediction of their model was that the rate of transcription elongation could control alternative splicing by determining the length of the window. Indeed, de la Mata and colleagues [17] showed that, in cultured human cells, a slow RNAPII promoted inclusion of the fibronectin EDI exon, and they proposed that slower transcription elongation expanded the “window of opportunity” for recognition of the weak 3’ splice site (3’SS) upstream of EDI before transcription of a competing 3’SS downstream. Evidence of kinetic coupling was obtained also in budding yeast by Howe *et al* [18], who observed enhanced second exon inclusion of modified *DYN2* transcripts in a slow RNAPII mutant or when cells were treated with chemical inhibitors of transcription. In striking contrast, slower elongation caused skipping of human CFTR exon 9 in a minigene construct due to enhanced recruitment of ETR-3, a negative splicing factor, to the 3’SS of the exon [19]. This demonstrates that slower transcription can also expand the window of opportunity for recruitment of negative factors that block splicing of a newly transcribed exon, reducing its inclusion rate. Overall, it seems that transcription rate determines the temporal window of opportunity for selection or rejection of an upstream sequence before a competing downstream sequence is transcribed. However, Fong et al [20] found that, in human cells, both faster and slower elongating RNA polymerase mutants may disrupt splicing in the same way, which seems contrary to the ‘‘window of opportunity’’ model. They proposed that an optimal rate of transcriptional elongation is required for normal co-transcriptional pre-mRNA splicing. Therefore, for the vast majority of genes, it is unclear what determines the splicing outcome of altering transcription elongation rate or how changes in transcription rate are regulated locally at alternatively spliced exons. It has been proposed that mechanisms may exist to slow or pause transcription downstream of introns in budding yeast, thereby stretching the window of opportunity for co-transcriptional spliceosome assembly to occur [6,21,22],

### Splicing fidelity

Splicing fidelity in yeast is the ability of the spliceosome to distinguish between optimal and suboptimal splice sites. Spliceosome assembly occurs in a stepwise fashion, and DExD/H-box RNA-stimulated NTPases are implicated in promoting fidelity at distinct steps by rejecting suboptimal substrates [23,24]. Mutations that reduce the NTPase activity of these factors were shown to increase the productive splicing of suboptimal substrates [23,25]. How transcription elongation rate affects splicing fidelity in budding yeast is an open question.

### Transcription rate affects splicing efficiency

To investigate the effect of transcription elongation rate on co-transcriptional splicing efficiency in budding yeast [26], we previously used RNAPII trigger loop mutants that elongate, on average, 4 times faster (Rpb1-*G1097D*) or 8 times slower (Rpb1-*H1085Y*) than WT RNAPII (~12 nt/s) *in vitro* [27]. Native Elongating Transcript (NET)-RTqPCR analysis for several genes showed that more splicing occurs co-transcriptionally with the slow mutant, and less with the fast mutant compared with wild-type RNAPII. This is compatible with both recruitment coupling and kinetic coupling; slower elongation allows more time for co-transcriptional recruitment of splicing factors to the nascent transcript, thereby promoting spliceosome assembly and splicing, whereas faster elongation has the opposite effect. Notably, analysis of total RNA revealed that overall splicing efficiency was also slightly improved with the slow RNAPII mutant and decreased with the fast mutant compared to wild-type. This indicates that splicing is more efficient when co-transcriptional, and that post-transcriptional splicing in the RNAPII fast mutant does not compensate for its reduced co-transcriptional splicing, which is, in itself, a possible raison d’etre for coupling transcription and splicing [26]. Interestingly, transcriptome-wide analysis of splicing efficiency by RNA sequencing revealed that the effect of the fast RNAPII mutant was mainly to reduce splicing efficiency with ribosomal protein (RP) coding transcripts, whereas the slow mutant increased the efficiency with both RP and non-RP transcripts.

### Transcription rate affects splicing fidelity

Using deep RNA sequencing, we previously measured splicing fidelity in RNAPII elongation mutants in which *UPF1* was deleted in order to protect mis-spliced transcripts from nonsense-mediated decay (NMD). We calculated the splicing error frequency (SEF) as a measure of splicing at “novel” versus annotated splice sites, where “novel” refers to splicing events on intron-containing transcripts that were not listed in the *Saccharomyces* annotation file (ENSEMBL, version R64-1-1.75). The SEF for each transcript is the number of reads that were aligned to novel splice junctions divided by the number of reads that were aligned to the annotated splice junction in that transcript. Notably, in the strain with WT RNAPII, non-RP transcripts showed, on average, 1000-fold higher SEF (frequent use of novel splice sites relative to annotated splice sites, i.e. low splicing fidelity) than RP transcripts. Considering the results from all three strains, WT as well as the RNAPII mutants, low SEF (higher splicing fidelity) correlated with longer introns together with more stable intron secondary structure, high 3’SS score and greater abundance, which are all features of RP transcripts [26].

## Materials and Methods

For details of the data generation and analysis methods, see [26].

## Results and Discussion

What makes RP transcripts more sensitive to faster transcription? An analysis of all RP introns, comparing splice site scores with splicing efficiency did not reveal a strong correlation overall. However, a closer inspection of the splicing efficiencies (Table S1), focusing on the 23 RP transcripts that have non-consensus 5’ splice sites (5’SS) (i.e. do not match the consensus sequence GUAUGU), reveals that the majority (18) were spliced less efficiently in the fast strain (Figure 1; Table S2). The most frequently occurring non-consensus 5’SS in the RP genes has the sequence GUACGU (Table S2), which Carrillo Oesterreich et al [2] showed, using a reporter gene, delays co-transcriptional splicing compared with a consensus 5’SS. Therefore, the reduced splicing efficiency observed for these RP transcripts in the fast mutant may be explained by the combination of delayed splicing due to the non-consensus 5’SS and the shorter window of opportunity for splicing to occur co-transcriptionally due to the faster transcription elongation. Consistent with this explanation is the observation that for 18 of the RP transcripts with non-consensus 5’SS, splicing was more efficient in the slow RNAPII strain, while one (*RPL22A*) was unaffected by the slow RNAPII mutation (Figure 1, Table S1). Moreover, *YML6*, which encodes a mitochondrial RP protein, also has 5’SS GUACGU and it too splices less efficiently in the fast and more efficiently in the slow RNAPII mutant.

**Figure 1.**
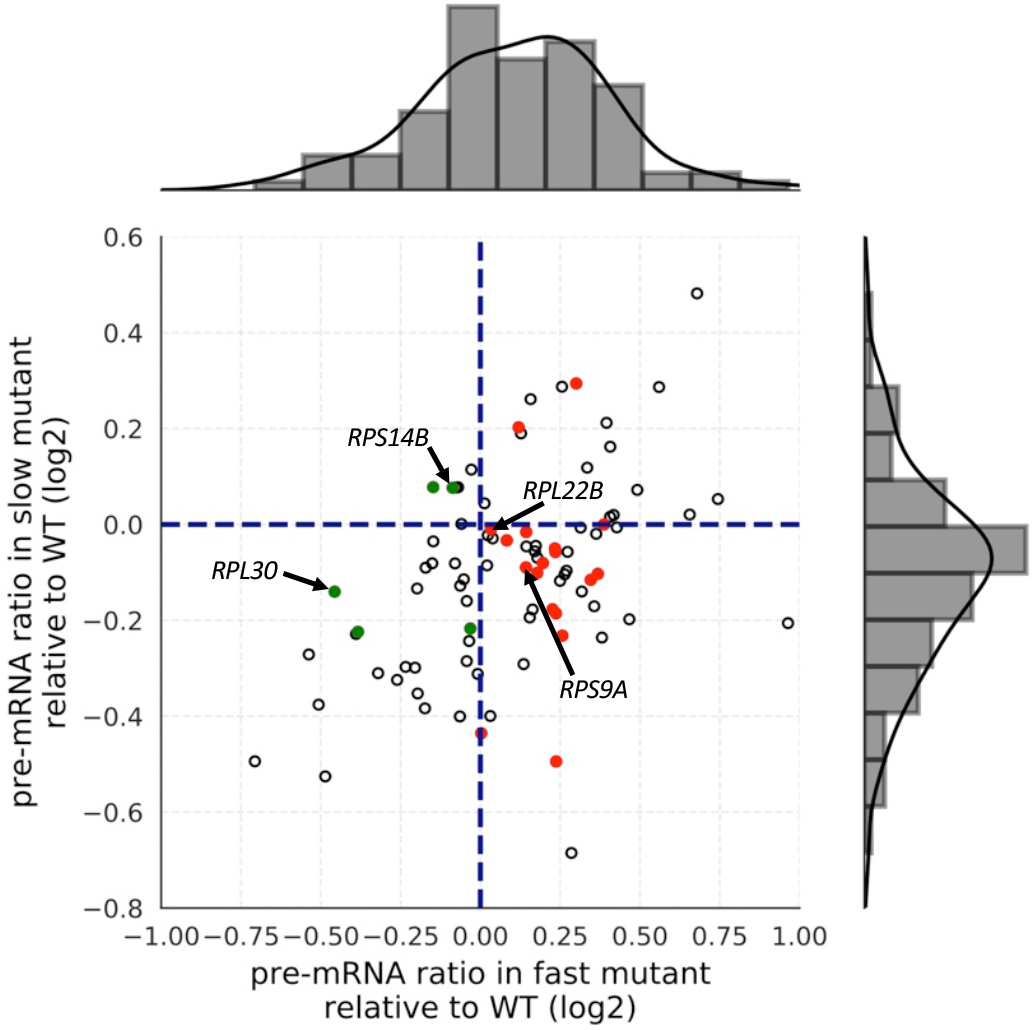
Faster transcription reduces and slower transcription increases the efficiency of splicing RP transcripts with non-canonical 5’ splice sites. Splicing efficiencies were calculated (Table S1) as RNA-seq read counts from pre-mRNA divided by total reads (pre-mRNA+mRNA) for each transcript [26], then the SEF for fast or slow RNAPII was divided by the corresponding SEF for the WT RNAPII and log2 values (columns Q and R in Table S1) were plotted. Transcripts that splice significantly more or less efficiently in the fast mutant compared to WT lie to the left and right of the vertical line respectively. Those that splice more or less efficiently in the slow mutant compared to WT lie below or above the horizontal line respectively. RP transcripts with non-consensus 5’SS are represented by green (more efficiently spliced in the fast mutant) or red (less efficiently spliced in the fast mutant) dots. Histograms show the distribution of pre-mRNA ratios relative to WT in the fast (top) and slow (right) mutants. In the fast mutant there are more genes with increased pre-mRNA ratio (reduced splicing efficiency) relative to WT and in the slow mutant there are more genes with reduced pre-mRNA ratio (increased splicing efficiency) relative to WT. Genes whose splicing is inhibited by their protein products are annoted in the figure.

For five RPs with non-consensus 5’SS (*RPS14B, RPS19A, RPS21B, RPL30* and *RPL43B*) splicing was more efficient in the fast RNAPII strain, suggesting that, for these transcripts, other factors have a greater influence on splicing efficiency than 5’SS. In the case of two of these transcripts, *RPS14B* and *RPL30*, their protein products Rps14p and Rpl30p, when in excess, bind to their respective precursor transcripts and inhibit splicing [28,29]. Therefore, in these two cases at least, competition between co-transcriptional spliceosome assembly and feedback inhibition of splicing by the protein product, may explain the different response to changes in transcription speed, for example if faster transcription through the intron permits spliceosome assembly or optimal secondary structure in the transcript before binding of the inhibitory protein. It would be interesting to investigate whether splicing of the other three transcripts in this category is also subject to negative regulation. On the other hand, *RPS9A* [30] and *RPL22B* [31] that have non-consensus 5’SS and whose splicing is inhibited by their protein products, are not spliced more efficiently with the fast RNAPII, so other factors come into play.

Curiously, the same trends are not apparent amongst non-RP transcripts with non-consensus 5’SS, but many of these have other atypical intron features that may affect their splicing kinetics. Nor is the same trend seen with RP transcripts that have suboptimal branchpoint sequences, although there are very few of these.

Next, the effect on splicing fidelity of changing transcription elongation rate was examined more closely, by comparing SEF scores in the RNAPII mutants with those in WT cells (P < 0.01 by Fisher’s Exact Test). Overall, there were more splicing events with reduced fidelity (100) than with increased fidelity (56) in the RNAPII mutants compared to WT, with RP transcripts being affected more than non-RP transcripts (105 and 51 respectively) (Fig 3 of [26]). We now note that, for 10 genes, fidelity of splicing at specific sites was increased by both fast and slow RNAPII mutants (Figure 2, green stars), while for a set of 24 distinct genes the fidelity of specific splicing events was decreased by both fast and slow RNAPII mutants (Figure 2, red stars). This observation cannot easily be explained by a window of opportunity model, and suggests that the length of time between transcription and splicing is not an important determinant of splicing fidelity, as might have been expected if co-transcriptional recruitment of fidelity factors was a determining step for splicing fidelity. The single exception was *RPL21B*, for which use of a novel 5’SS, 79 bases downstream of the canonical 5’SS, occurred with reduced frequency in the slow RNAPII mutant and with increased frequency in the fast RNAPII mutant compared with WT. Overall, for both RP and non-RP genes, the effect on splicing fidelity of changing transcription speed (either faster or slower) seems to be gene and splice site specific.

**Figure 2.**
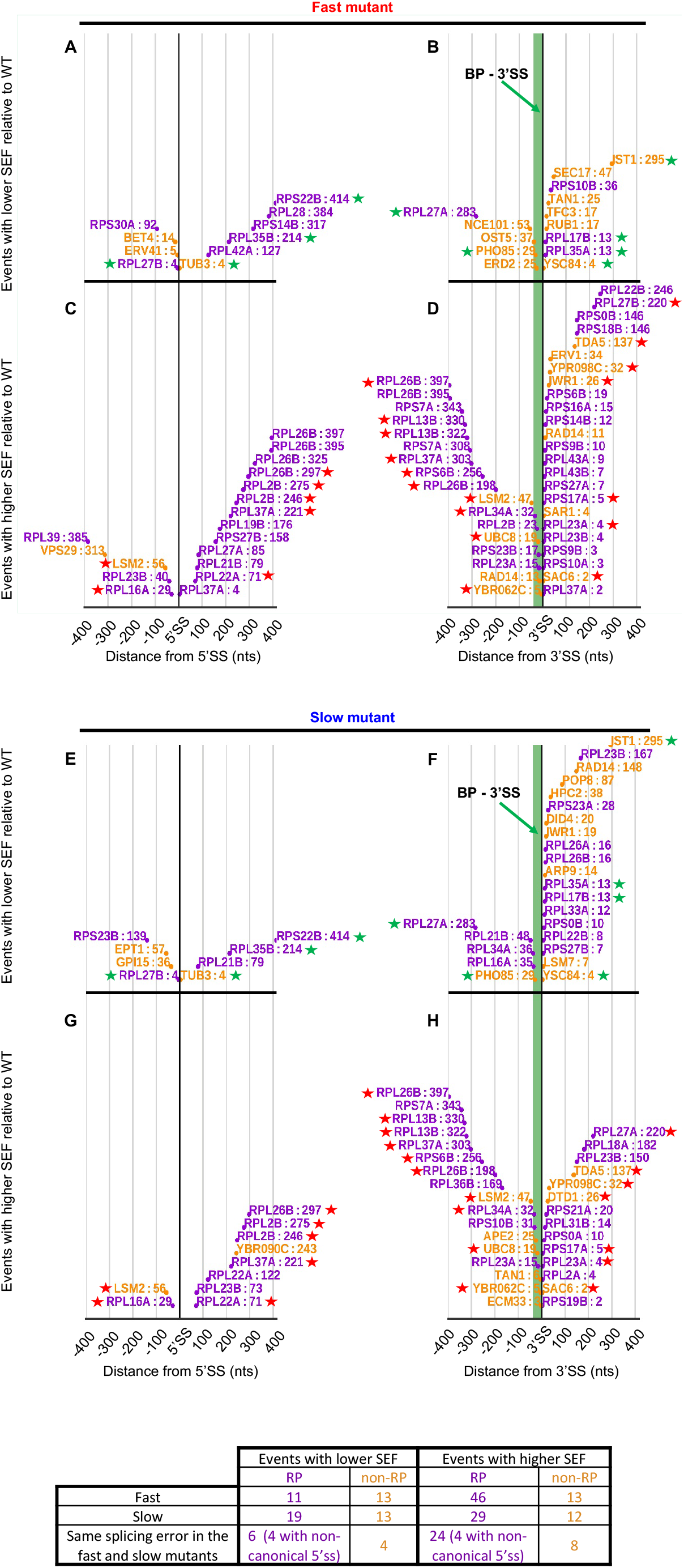
Distribution of the novel splice sites in RP (purple) and non-RP (orange) intron-containing transcripts whose SEF is significantly different in the fast and slow mutants relative to WT (P < 0.01). (A,B) Alternative 5’SS and 3’SS splicing events with increased fidelity in the fast mutant. (C,D) Alternative 5’SS and 3’SS splicing events with reduced fidelity in the fast mutant. (E,F) Alternative 5’SS and 3’SS splicing events with increased fidelity in the slow mutant. (G,H) Alternative 5’SS and 3’SS splicing events with reduced fidelity in the slow mutant. Green region upstream of 3’SS is the mean distance between BP and annotated 3’SS in budding yeast (37 nt). Stars indicate splicing events that have SEF significantly different to WT in the same direction-either lower SEF (green) or higher SEF (red) - in **both** fast and slow mutants. Modified from Figure 3 in [26]. The table below states the number of splicing events for RP versus nonRP transcripts in the fast versus slow mutants with increased or decreased SEF.

How can this be explained? Changing the elongation rate may alter the local chromatin environment and multiple properties of nascent RNA in a gene-specific manner, affecting the availability of cryptic splice sites in yeast transcripts, as has been suggested to explain the effects of transcription speed on alternative splicing events [13,32,33]. Alternatively, it is conceivable that NTP-dependent kinetic proofreading mechanisms in yeast require an optimum transcription speed for each splicing event, that is fine-tuned to the rate of NTP hydrolysis [24,34].

### Concluding remarks

We previously reported that RP transcripts are spliced faster and more co-transcriptionally [35,36] and that they tend to be spliced with greater efficiency and higher fidelity [26]. Additionally, compared with non-RP transcripts, the splicing efficiency and fidelity of RP transcripts is more sensitive to changes in transcription elongation rate [26]. Taken together, these results indicate that the splicing of RP transcripts is more functionally coupled to transcription than that of non-RP transcripts, and that this coupling is highly beneficial. This supports a proposal based on modelling [8], that co-transcriptional splicing is more efficient than post-transcriptional splicing. Yet, it remains largely unknown what determines the extent to which splicing of a transcript is more or less coupled to transcription.

Our observation that a non-consensus 5’SS confers sensitivity to transcription elongation rate supports both the recruitment and kinetic coupling models, and suggests that, at least for RP transcripts, a non-consensus 5’SS may reduce the efficiency of spliceosome assembly and/or splicing catalysis in a manner that is compensated by slower transcription elongation. This is compatible with the observation of Carrillo Oesterreich et al [2] that changing the GUAUGU consensus 5’SS of an RP intron in a reporter transcript to GUACGU, delays co-transcriptional splicing, which was proposed to be due to weakened 5’SS base-pairing with U1 snRNA delaying spliceosome assembly [2]. There are several reports of U1 snRNAs interacting with components of the transcription machinery [37–41], suggesting a possible role for U1 snRNPs in coupling transcription and splicing. In this senario, recruitment coupling of U1 snRNPs by the transcription machinery could be particularly beneficial for efficient splicing of RP transcripts with non-consensus 5’SSs, and might explain the observed sensitivity of splicing efficiency to changes in transcription speed.

However, it is questionable whether replacing U with C as the base at position +4 of a 5’SS alters the stability of the 5’SS:U1 snRNA interaction, as the opposing base at position +5 in U1 snRNA is pseudoU [42], a modified base that can pair with either U or C [43]. Indeed, in the study by Carrillo Oesterreich et al [2]; U1 snRNP was apparently recruited equally efficiently to reporter transcripts containing either GUAUGU or GUACGU at the 5’SS, whereas the U1 snRNP signal was lost more slowly from the GUACGU reporter. A possible explanation for this delay is the reduced ability of CGU in the nonconsensus 5’SS to pair with ACA in the ACAGAGA motif of U6 snRNA, which is coupled with U1 snRNP displacement from the 5’SS by Prp28 [44–46]. In this case, slower transcription elongation would allow more time for the assembling spliceosome to proceed through this checkpoint and/or for co-transcriptional recruitment of factors required to form catalytically active spliceosomes.

Genome-wide studies in mammalian cells also found that specific transcript features, such as suboptimal splice sites, may make a particular splicing event more sensitive to transcription rate, especially in transcripts that encode RNA binding or RNA processing factors [20,47]. Splicing fidelity could be considered to be a form of alternative splicing. Indeed, a human cell line carrying a slow RNAPII mutant was reported to increase fidelity for inclusion of a subset of alternatively splicing exons while, at the same time, reducing inclusion of another subset, giving rise to the proposal that there is an optimal transcription elongation rate for individual splicing events [20].

Synthesis of the large number of ribosomal proteins is metabolically expensive for cells, and their expression is tightly regulated. As many RP genes in budding yeast contain introns, it is logical that the splicing of their transcripts is also highly efficient and accurate. However, high splicing fidelity can constrain evolution by reducing the chance of generating new transcript or protein isoforms from the same gene [48]. It has been proposed that alternative splicing is a source of functional innovation in higher eukaryotes and a mechanism for evolutionary diversification of cell types [48,49]. Conceivably, the higher rate of splicing errors in non-RP transcripts reflects reduced constraints, allowing alternative splicing events that offer evolutionary advantage, such as under different environmental conditions. For example, in response to heat shock, the *APE2* transcript in budding yeast is spliced to produce a mRNA 18 nts longer than the annotated mRNA by using an upstream alternative 3’SS [50]; likely an evolutionary adaptation to heat stress. Therefore, the coupling of transcription and splicing may be more advantageous to RP genes, which function in key metabolic processes and whose regulation is highly conserved, whereas non-RP genes escape these constraints.

What other features of RP transcripts cause their splicing to be more coupled to transcription? Promotor structure was shown to be important for alternative splicing in human cells [51] and yeast [52], and it was shown that RP transcripts in budding yeast have distinct promoter architectures, with an exclusive pattern of DNA binding proteins that enhance their transcription [53]. One informative future approach would be to study the effect on co-transcriptional splicing of swapping the promoters and different intron features of yeast RP and non-RP transcripts.

## Supporting information

## Acknowledgements

We are grateful to Edward Wallace for helpful comments about the manuscript. This work was supported by Wellcome funding to JB [104648] and a Darwin Trust of Edinburgh studentship to VA. Work in the Wellcome Centre for Cell Biology is supported by Wellcome core funding [092076].

## References

[1] Ameur A, Zaghlool A, Halvardson J, et al. Total RNA sequencing reveals nascent transcription and widespread co-transcriptional splicing in the human brain. Nat Struct Mol Biol. 2011;18:1435–1440.

[2] Carrillo Oesterreich F, Herzel L, Straube K, et al. Splicing of Nascent RNA Coincides with Intron Exit from RNA Polymerase II. Cell. 2016;165:372–381.

[3] Herzel L, Straube K, Neugebauer KM. Long-read sequencing of nascent RNA reveals coupling among RNA processing events. Genome Res. 2018;28:1008–1019.

[4] Khodor YL, Rodriguez J, Abruzzi KC, et al. Nascent-seq indicates widespread cotranscriptional pre-mRNA splicing in Drosophila. Genes Dev. 2011;25:2502–2512.

[5] Listerman I, Sapra AK, Neugebauer KM. Cotranscriptional coupling of splicing factor recruitment and precursor messenger RNA splicing in mammalian cells. Nat Struct Mol Biol. 2006;13:815–822.

[6] Carrillo Oesterreich F, Preibisch S, Neugebauer KM. Global Analysis of Nascent RNA Reveals Transcriptional Pausing in Terminal Exons. Molecular Cell. 2010;40:571–581.

[7] Tilgner H, Knowles DG, Johnson R, et al. Deep sequencing of subcellular RNA fractions shows splicing to be predominantly co-transcriptional in the human genome but inefficient for lncRNAs. Genome Res. 2012;22:1616–1625.

[8] Aitken S, Alexander RD, Beggs JD. Modelling Reveals Kinetic Advantages of Co-Transcriptional Splicing. PLOS Comput Biol. 2011;7:e1002215.

[9] Li X, Manley JL. Cotranscriptional processes and their influence on genome stability. Genes Dev. 2006;20:1838–1847.

[10] Naftelberg S, Schor IE, Ast G, et al. Regulation of Alternative Splicing Through Coupling with Transcription and Chromatin Structure. Annual Review of Biochemistry. 2015;84:165–198.

[11] Skourti-Stathaki K, Proudfoot NJ. A double-edged sword: R loops as threats to genome integrity and powerful regulators of gene expression. Genes Dev. 2014;28:1384–1396.

[12] Moehle EA, Braberg H, Krogan NJ, et al. Adventures in time and space. RNA Biology. 2014;11:313–319.

[13] Dujardin G, Lafaille C, Petrillo E, et al. Transcriptional elongation and alternative splicing. Biochimica et Biophysica Acta (BBA) - Gene Regulatory Mechanisms. 2013;1829:134–140.

[14] Bentley DL. Coupling mRNA processing with transcription in time and space. Nat Rev Genet. 2014;15:163–175.

[15] Kornblihtt AR, Schor IE, Alló M, et al. Alternative splicing: a pivotal step between eukaryotic transcription and translation. Nat Rev Mol Cell Biol. 2013;14:153–165.

[16] Eperon LP, Graham IR, Griffiths AD, et al. Effects of RNA secondary structure on alternative splicing of pre-mRNA: is folding limited to a region behind the transcribing RNA polymerase? Cell. 1988;54:393–401.

[17] de la Mata M, Alonso CR, Kadener S, et al. A Slow RNA Polymerase II Affects Alternative Splicing In Vivo. Molecular Cell. 2003;12:525–532.

[18] Howe KJ, Kane CM, Ares M. Perturbation of transcription elongation influences the fidelity of internal exon inclusion in Saccharomyces cerevisiae. RNA. 2003;9:993–1006.

[19] Dujardin G, Lafaille C, de la Mata M, et al. How Slow RNA Polymerase II Elongation Favors Alternative Exon Skipping. Molecular Cell. 2014;54:683–690.

[20] Fong N, Kim H, Zhou Y, et al. Pre-mRNA splicing is facilitated by an optimal RNA polymerase II elongation rate. Genes Dev. 2014;28:2663–2676.

[21] Alexander RD, Innocente SA, Barrass JD, et al. Splicing-Dependent RNA Polymerase Pausing in Yeast. Molecular Cell. 2010;40:582–593.

[22] Milligan L, Sayou C, Tuck A, et al. RNA polymerase II stalling at pre-mRNA splice sites is enforced by ubiquitination of the catalytic subunit. Proudfoot NJ, editor. eLife. 2017;6:e27082.

[23] Query CC, Konarska MM. Splicing fidelity revisited. Nat Struct Mol Biol. 2006;13:472–474.

[24] Semlow DR, Staley JP. Staying on message: ensuring fidelity in pre-mRNA splicing. Trends in Biochemical Sciences. 2012;37:263–273.

[25] Burgess SM, Guthrie C. A mechanism to enhance mRNA splicing fidelity: The RNA-dependent ATPase Prp16 governs usage of a discard pathway for aberrant lariat intermediates. Cell. 1993;73:1377–1391.

[26] Aslanzadeh V, Huang Y, Sanguinetti G, et al. Transcription rate strongly affects splicing fidelity and cotranscriptionality in budding yeast. Genome Res. 2018;28:203–213.

[27] Kaplan CD, Jin H, Zhang IL, et al. Dissection of Pol II Trigger Loop Function and Pol II Activity–Dependent Control of Start Site Selection In Vivo. PLoS Genet. 2012;8:e1002627.

[28] Eng FJ, Warner JR. Structural basis for the regulation of splicing of a yeast messenger RNA. Cell. 1991;65:797–804.

[29] Fewell SW, Woolford JL. Ribosomal protein S14 of Saccharomyces cerevisiae regulates its expression by binding to RPS14B pre-mRNA and to 18S rRNA. Mol. Cell. Biol. 1999;19:826–834.

[30] Petibon C, Parenteau J, Catala M, et al. Introns regulate the production of ribosomal proteins by modulating splicing of duplicated ribosomal protein genes. Nucleic Acids Res. 2016;44:3878–3891.

[31] Gabunilas J, Chanfreau G. Splicing-Mediated Autoregulation Modulates Rpl22p Expression in Saccharomyces cerevisiae. PLOS Genetics. 2016;12:e1005999.

[32] Buratti E, Baralle FE. Influence of RNA Secondary Structure on the Pre-mRNA Splicing Process. Mol. Cell. Biol. 2004;24:10505–10514.

[33] Saldi T, Cortazar MA, Sheridan RM, et al. Coupling of RNA Polymerase II Transcription Elongation with Pre-mRNA Splicing. Journal of Molecular Biology. 2016;428:2623–2635.

[34] Hopfield JJ. Kinetic Proofreading: A New Mechanism for Reducing Errors in Biosynthetic Processes Requiring High Specificity. PNAS. 1974;71:4135–4139.

[35] Barrass JD, Reid JEA, Huang Y, et al. Transcriptome-wide RNA processing kinetics revealed using extremely short 4tU labeling. Genome Biology. 2015;16:282.

[36] Wallace EWJ, Beggs JD. Extremely fast and incredibly close: co-transcriptional splicing in budding yeast. RNA. 2017;rna.060830.117.

[37] Phatnani HP, Greenleaf AL. Phosphorylation and functions of the RNA polymerase II CTD. Genes Dev. 2006;20:2922–2936.

[38] Damgaard CK, Kahns S, Lykke-Andersen S, et al. A 5’ splice site enhances the recruitment of basal transcription initiation factors in vivo. Mol. Cell. 2008;29:271–278.

[39] Fong YW, Zhou Q. Stimulatory effect of splicing factors on transcriptional elongation. Nature. 2001;414:929–933.

[40] Das R, Yu J, Zhang Z, et al. SR Proteins Function in Coupling RNAP II Transcription to Pre-mRNA Splicing. Molecular Cell. 2007;26:867–881.

[41] Spiluttini B, Gu B, Belagal P, et al. Splicing-independent recruitment of U1 snRNP to a transcription unit in living cells. J. Cell. Sci. 2010;123:2085–2093.

[42] Plaschka C, Lin P-C, Charenton C, et al. Prespliceosome structure provides insights into spliceosome assembly and regulation. Nature. 2018;559:419.

[43] Kierzek E, Malgowska M, Lisowiec J, et al. The contribution of pseudouridine to stabilities and structure of RNAs. Nucleic Acids Res. 2014;42:3492–3501.

[44] Staley JP, Guthrie C. An RNA switch at the 5’ splice site requires ATP and the DEAD box protein Prp28p. Mol. Cell. 1999;3:55–64.

[45] Chen JY-F, Stands L, Staley JP, et al. Specific Alterations of U1-C Protein or U1 Small Nuclear RNA Can Eliminate the Requirement of Prp28p, an Essential DEAD Box Splicing Factor. Molecular Cell. 2001;7:227–232.

[46] Kuhn AN, Li Z, Brow DA. Splicing Factor Prp8 Governs U4/U6 RNA Unwinding during Activation of the Spliceosome. Molecular Cell. 1999;3:65–75.

[47] Ip JY, Schmidt D, Pan Q, et al. Global impact of RNA polymerase II elongation inhibition on alternative splicing regulation. Genome Res. 2011;21:390–401.

[48] Keren H, Lev-Maor G, Ast G. Alternative splicing and evolution: diversification, exon definition and function. Nature Reviews Genetics. 2010;11:345–355.

[49] Bush SJ, Chen L, Tovar-Corona JM, et al. Alternative splicing and the evolution of phenotypic novelty. Philos Trans R Soc Lond B Biol Sci. 2017;372.

[50] Meyer M, Plass M, Pérez-Valle J, et al. Deciphering 3’ss Selection in the Yeast Genome Reveals an RNA Thermosensor that Mediates Alternative Splicing. Molecular Cell. 2011;43:1033–1039.

[51] Cramer P, Pesce CG, Baralle FE, et al. Functional association between promoter structure and transcript alternative splicing. Proc Natl Acad Sci U S A. 1997;94:11456–11460.

[52] Kawashima T, Douglass S, Gabunilas J, et al. Widespread Use of Non-productive Alternative Splice Sites in Saccharomyces cerevisiae. PLoS Genet. 2014;10:e1004249.

[53] Knight B, Kubik S, Ghosh B, et al. Two distinct promoter architectures centered on dynamic nucleosomes control ribosomal protein gene transcription. Genes Dev. 2014;28:1695–1709.

