## Supplementary material for "Revisiting the window of opportunity for co-transcriptional splicing efficiency and fidelity"

**Supplemental Information**

**Table S1**

Sheet 1: RNA-seq read counts for pre-mRNAs in the fast RNAPII mutant (columns A,B show counts for 2 biological replicates), slow mutant (C,D), wild-type (WT; E,F) or for spliced mRNAs (H-M). Pre-mRNA read counts calculated as sum of the reads aligning to 5'SS, intron, 3'SS and the reads spanning on exon1-intron-exon2 sequence. mRNA read counts calculated as sum of the reads aligning to exon1 – exon2 splice junction. Pre-mRNA ratios for fast, slow and WT (columns N,O,P respectively were calculated as counts of pre-mRNA divided by total read counts (pre-mRNA+mRNA)); log2 ratio of mutant/WT for fast (S) and slow (T) RNAPII mutants.

Sheet 2 (named 'RP'): Data selected from sheet 1 for RP transcripts only, indicating those with non-consensus 5'SS that splice more efficiently (green) or less efficiently (red) in the fast RNAPII mutant.

| | premRNA_fast_A | premRNA_fast_B | premRNA_slow_A | premRNA_slow_B | premRNA_wt_A | premRNA_wt_B | mRNA_fast_A | mRNA_fast_B | mRNA_slow_A | mRNA_slow_B | mRNA_wt_A | mRNA_wt_B | premRNA ratio_fast | premRNA ratio_slow | premRNA ratio_wt | premRNAratio_fast/wt (log2) | premRNA ratio_slow/wt(log2) |
|---|---|---|---|---|---|---|---|---|---|---|---|---|---|---|---|---|---|
| YLR445W | 22 | 63 | 38 | 61 | 17 | 22 | 0 | 30 | 2 | 9 | 25 | 5 | 0.83871 | 0.910714 | 0.609788 | 0.459863 | 0.57869 |
| RPS21A | 2681 | 2194 | 1338 | 2088 | 2601 | 2657 | 31967 | 32994 | 19657 | 27747 | 29812 | 27827 | 0.069865 | 0.066857 | 0.083703 | -0.26072 | -0.3242 |
| RPS21B | 886 | 935 | 744 | 1058 | 1233 | 1217 | 28754 | 29480 | 20819 | 30790 | 31625 | 28061 | 0.030317 | 0.033862 | 0.039546 | -0.38342 | -0.22387 |
| YBL059W | 340 | 526 | 291 | 445 | 298 | 371 | 646 | 442 | 390 | 631 | 243 | 272 | 0.444108 | 0.420441 | 0.563907 | -0.34455 | -0.42356 |
| RAD14 | 86 | 150 | 135 | 110 | 47 | 55 | 246 | 289 | 300 | 294 | 187 | 225 | 0.300361 | 0.291311 | 0.198642 | 0.596529 | 0.552392 |
| COF1 | 922 | 1075 | 644 | 1268 | 958 | 1017 | 5859 | 5744 | 6647 | 9631 | 6876 | 6690 | 0.146808 | 0.102335 | 0.127123 | 0.207708 | -0.31293 |
| YBR090C | 1265 | 1073 | 1472 | 1733 | 1638 | 1438 | 2033 | 1762 | 2070 | 2534 | 1831 | 1523 | 0.381025 | 0.410862 | 0.478914 | -0.32988 | -0.22111 |
| LSB3 | 197 | 172 | 142 | 238 | 292 | 213 | 1737 | 1439 | 2112 | 2254 | 1729 | 1546 | 0.104314 | 0.079252 | 0.132787 | -0.34819 | -0.74459 |
| RPS11B | 2947 | 3426 | 1945 | 2912 | 3469 | 2996 | 23216 | 21589 | 19059 | 23283 | 29458 | 25085 | 0.124799 | 0.101884 | 0.106023 | 0.235231 | -0.05745 |
| RPS11A | 2284 | 2349 | 1378 | 2189 | 2451 | 2033 | 44844 | 40867 | 41816 | 49592 | 60786 | 47667 | 0.051409 | 0.037088 | 0.039832 | 0.368095 | -0.10297 |
| YDR261W- | 13 | 10 | 1 | 87 | 13 | 9 | 211 | 165 | 203 | 318 | 128 | 136 | 0.057589 | 0.109858 | 0.077134 | -0.42156 | 0.51021 |
| YIL156W-B | 347 | 214 | 429 | 621 | 322 | 376 | 599 | 860 | 777 | 1057 | 846 | 965 | 0.283031 | 0.362902 | 0.278036 | 0.025688 | 0.384308 |
| IWR1 | 725 | 1321 | 967 | 1737 | 1383 | 1288 | 311 | 372 | 351 | 381 | 124 | 244 | 0.740039 | 0.7769 | 0.879224 | -0.24863 | -0.1785 |
| CIN2 | 776 | 750 | 532 | 808 | 724 | 556 | 206 | 280 | 197 | 176 | 218 | 205 | 0.75919 | 0.775453 | 0.749598 | 0.018344 | 0.048922 |
| NCB2 | 163 | 106 | 112 | 243 | 255 | 209 | 404 | 484 | 530 | 597 | 310 | 431 | 0.233569 | 0.23187 | 0.388945 | -0.73571 | -0.74625 |
| HRB1 | 1049 | 1507 | 1359 | 2059 | 1902 | 1896 | 447 | 448 | 614 | 702 | 563 | 371 | 0.736024 | 0.717272 | 0.803975 | -0.1274 | -0.16463 |
| GPI15 | 111 | 149 | 167 | 228 | 189 | 191 | 325 | 569 | 495 | 801 | 595 | 460 | 0.231054 | 0.23692 | 0.267233 | -0.20987 | -0.1737 |
| MOB1 | 151 | 174 | 242 | 386 | 238 | 201 | 251 | 305 | 262 | 461 | 232 | 271 | 0.369439 | 0.467942 | 0.466115 | -0.33535 | 0.005644 |
| CPT1 | 51 | 71 | 42 | 152 | 115 | 81 | 1334 | 1662 | 1295 | 1843 | 1484 | 1566 | 0.038896 | 0.053802 | 0.06055 | -0.6385 | -0.17047 |
| RPS19B | 2845 | 2911 | 1683 | 2603 | 2719 | 2406 | 25124 | 25245 | 22186 | 28165 | 27950 | 25344 | 0.102554 | 0.077555 | 0.08768 | 0.226072 | -0.17701 |
| RPS19A | 2397 | 2393 | 1629 | 2372 | 2452 | 2225 | 16794 | 16373 | 13905 | 17881 | 15652 | 15929 | 0.12621 | 0.110993 | 0.129001 | -0.03156 | -0.21692 |
| BOS1 | 246 | 293 | 220 | 185 | 211 | 268 | 225 | 223 | 272 | 389 | 224 | 255 | 0.545061 | 0.384727 | 0.498743 | 0.128122 | -0.37446 |
| YLR464W | 405 | 506 | 369 | 533 | 472 | 428 | 0 | 0 | 0 | 0 | 0 | 0 | 1 | 1 | 1 | 0 | 0 |
| ACT1 | 797 | 591 | 623 | 1124 | 762 | 888 | 9313 | 9012 | 9679 | 14803 | 12787 | 11723 | 0.070188 | 0.065523 | 0.063328 | 0.148393 | 0.049165 |
| RPS6B | 4319 | 4714 | 2009 | 3067 | 3773 | 3055 | 13989 | 16458 | 11840 | 18193 | 17903 | 16307 | 0.22928 | 0.144663 | 0.165923 | 0.466594 | -0.19782 |
| RPS6A | 1627 | 1788 | 1264 | 2138 | 1999 | 1861 | 6254 | 6308 | 6391 | 9254 | 8148 | 8710 | 0.213648 | 0.176398 | 0.186526 | 0.195859 | -0.08054 |
| RPO26 | 388 | 518 | 656 | 800 | 774 | 761 | 1785 | 1731 | 2349 | 2777 | 2945 | 2348 | 0.20444 | 0.220977 | 0.226447 | -0.1475 | -0.03528 |
| MEI4 | 44 | 119 | 54 | 99 | 91 | 193 | 0 | 0 | 10 | 0 | 0 | 0 | 1 | 0.921875 | 1 | 0 | -0.11736 |
| DMC1 | 547 | 662 | 268 | 597 | 375 | 374 | 4 | 32 | 0 | 42 | 47 | 8 | 0.973315 | 0.967136 | 0.933842 | 0.05973 | 0.050541 |
| APE2 | 3865 | 3726 | 3326 | 5183 | 3850 | 3673 | 535 | 639 | 440 | 541 | 419 | 381 | 0.866009 | 0.894325 | 0.903935 | -0.06184 | -0.01542 |
| SPT14 | 448 | 529 | 308 | 381 | 522 | 290 | 55 | 71 | 31 | 34 | 58 | 52 | 0.886161 | 0.913313 | 0.873977 | 0.019975 | 0.063515 |
| RPS22B | 4582 | 4571 | 2017 | 3779 | 5328 | 5360 | 14320 | 13836 | 8298 | 9625 | 11326 | 9231 | 0.245369 | 0.238736 | 0.343636 | -0.48593 | -0.52547 |
| RPL42B | 2253 | 2601 | 1548 | 2454 | 2372 | 2549 | 14555 | 13001 | 8923 | 11540 | 16536 | 15128 | 0.150376 | 0.161599 | 0.134824 | 0.157499 | 0.261339 |
| TDA5 | 122 | 171 | 135 | 199 | 367 | 305 | 404 | 354 | 447 | 521 | 297 | 260 | 0.278827 | 0.254174 | 0.546267 | -0.97024 | -1.10379 |
| KIN28 | 425 | 330 | 508 | 670 | 460 | 374 | 297 | 232 | 228 | 247 | 169 | 189 | 0.587916 | 0.71043 | 0.697809 | -0.24722 | 0.025861 |
| BET1 | 113 | 235 | 173 | 398 | 250 | 246 | 448 | 563 | 532 | 1001 | 676 | 518 | 0.247956 | 0.264939 | 0.295984 | -0.25543 | -0.15986 |
| ARP9 | 82 | 92 | 139 | 333 | 137 | 107 | 557 | 528 | 719 | 736 | 461 | 487 | 0.138356 | 0.236755 | 0.204616 | -0.56453 | 0.210479 |
| RPS10A | 2958 | 3108 | 2017 | 3432 | 3742 | 3886 | 39842 | 37670 | 35960 | 43875 | 45573 | 40250 | 0.072665 | 0.062829 | 0.081963 | -0.17371 | -0.38353 |
| RPS10B | 2235 | 2413 | 1848 | 2647 | 2051 | 2076 | 30938 | 27425 | 24684 | 28147 | 24466 | 21717 | 0.074122 | 0.077805 | 0.0823 | -0.15098 | -0.08102 |
| TMA20 | 286 | 360 | 160 | 412 | 313 | 254 | 830 | 1032 | 1250 | 1677 | 1066 | 1376 | 0.257447 | 0.155349 | 0.191402 | 0.427666 | -0.30109 |
| YDR535C | 643 | 465 | 354 | 315 | 175 | 289 | 0 | 0 | 0 | 0 | 0 | 0 | 1 | 1 | 1 | 0 | 0 |

| | premRNA_fast_A | premRNA_fast_B | premRNA_slow_A | premRNA_slow_B | premRNA_wt_A | premRNA_wt_B | mRNA_fast_A | mRNA_fast_B | mRNA_slow_A | mRNA_slow_B | mRNA_wt_A | mRNA_wt_B | premRNA ratio_fast | premRNA ratio_slow | premRNA ratio_wt | premRNAratio_fast/wt (log2) | premRNA ratio_slow/wt(log2) |
|---|---|---|---|---|---|---|---|---|---|---|---|---|---|---|---|---|---|
| RPL42A | 11151 | 11837 | 12080 | 16110 | 16205 | 16179 | 5470 | 4413 | 4676 | 5711 | 6610 | 5455 | 0.699665 | 0.729608 | 0.729064 | -0.05938 | 0.001075 |
| ARP2 | 226 | 208 | 208 | 318 | 243 | 205 | 1577 | 1737 | 2277 | 3003 | 2128 | 1711 | 0.116144 | 0.089728 | 0.104741 | 0.149084 | -0.22319 |
| UBC8 | 118 | 126 | 168 | 173 | 104 | 82 | 153 | 184 | 164 | 361 | 111 | 158 | 0.420938 | 0.414997 | 0.412694 | 0.028536 | 0.008029 |
| UBC9 | 200 | 203 | 190 | 270 | 217 | 275 | 1324 | 1333 | 1019 | 1496 | 1456 | 1286 | 0.131698 | 0.155021 | 0.152938 | -0.21572 | 0.019518 |
| HPC2 | 29 | 111 | 47 | 136 | 94 | 95 | 183 | 147 | 187 | 222 | 103 | 150 | 0.283513 | 0.290371 | 0.432456 | -0.60914 | -0.57465 |
| MAF1 | 115 | 129 | 87 | 146 | 114 | 163 | 502 | 558 | 276 | 543 | 386 | 366 | 0.187079 | 0.225785 | 0.268064 | -0.51893 | -0.24763 |
| YRA1 | 6291 | 6815 | 4152 | 6033 | 3628 | 3337 | 3977 | 3961 | 2845 | 3630 | 2571 | 2972 | 0.622552 | 0.608869 | 0.557091 | 0.160281 | 0.128217 |
| ASC1 | 4257 | 4757 | 2087 | 3579 | 3332 | 3246 | 34037 | 38177 | 21126 | 34035 | 33712 | 36569 | 0.110982 | 0.092529 | 0.085737 | 0.372336 | 0.109981 |
| RPS18A | 2935 | 2986 | 2103 | 3215 | 3425 | 3251 | 29420 | 29306 | 23611 | 33687 | 30949 | 33262 | 0.091591 | 0.084453 | 0.094338 | -0.04264 | -0.15968 |
| RPS18B | 1113 | 1204 | 971 | 1151 | 1045 | 1252 | 23529 | 24150 | 21514 | 31723 | 30896 | 32434 | 0.046327 | 0.039098 | 0.034942 | 0.40691 | 0.162161 |
| PBA1 | 96 | 153 | 99 | 193 | 147 | 150 | 187 | 286 | 288 | 351 | 211 | 232 | 0.343871 | 0.305297 | 0.401642 | -0.22404 | -0.3957 |
| YLR211C | 12 | 11 | 34 | 84 | 37 | 27 | 34 | 86 | 102 | 129 | 47 | 38 | 0.187136 | 0.322183 | 0.42793 | -1.19329 | -0.4095 |
| MRPL44 | 233 | 270 | 418 | 595 | 421 | 334 | 1146 | 1178 | 2220 | 2613 | 760 | 1017 | 0.177714 | 0.171964 | 0.301851 | -0.76428 | -0.81173 |
| RPL14A | 4015 | 4266 | 3099 | 3925 | 4950 | 4539 | 70593 | 66837 | 46404 | 65645 | 70143 | 65670 | 0.056906 | 0.05951 | 0.065284 | -0.19815 | -0.1336 |
| RPL31A | 2001 | 2007 | 1469 | 2027 | 1901 | 1935 | 61377 | 58478 | 41713 | 51267 | 58439 | 50652 | 0.032377 | 0.036027 | 0.03415 | -0.07693 | 0.077154 |
| RPL31B | 947 | 1042 | 993 | 1240 | 1704 | 1264 | 9305 | 9366 | 11632 | 15559 | 13204 | 13276 | 0.096244 | 0.076234 | 0.100617 | -0.06411 | -0.40037 |
| RPL14B | 2369 | 2245 | 1641 | 2333 | 2949 | 2462 | 24791 | 23821 | 14367 | 19682 | 18894 | 18656 | 0.086676 | 0.104242 | 0.125796 | -0.53739 | -0.27115 |
| ERD2 | 205 | 146 | 148 | 211 | 120 | 94 | 454 | 488 | 394 | 862 | 558 | 444 | 0.270681 | 0.234854 | 0.175856 | 0.622196 | 0.417367 |
| RPL37B | 2912 | 3477 | 1721 | 2196 | 1860 | 1567 | 13952 | 11459 | 8860 | 10506 | 11712 | 9786 | 0.202734 | 0.167768 | 0.137536 | 0.559781 | 0.286659 |
| RPL16B | 1573 | 1746 | 1120 | 1453 | 1863 | 1484 | 28001 | 27449 | 26926 | 33026 | 36431 | 35372 | 0.056497 | 0.041038 | 0.044457 | 0.345744 | -0.11546 |
| RPL16A | 2026 | 2146 | 1419 | 2041 | 2529 | 2198 | 17526 | 16226 | 17937 | 22324 | 24017 | 22443 | 0.110215 | 0.078539 | 0.092235 | 0.256934 | -0.2319 |
| RPL37A | 3353 | 3119 | 2428 | 3464 | 3589 | 3342 | 7605 | 7058 | 6854 | 7806 | 8238 | 7791 | 0.306231 | 0.284473 | 0.301823 | 0.020915 | -0.08541 |
| RPS17A | 8545 | 7274 | 6069 | 7438 | 9703 | 8882 | 17861 | 17134 | 15351 | 20995 | 26661 | 21157 | 0.310809 | 0.272465 | 0.281256 | 0.144144 | -0.04581 |
| RFA2 | 134 | 192 | 133 | 208 | 170 | 124 | 320 | 333 | 233 | 390 | 374 | 298 | 0.330434 | 0.355607 | 0.303169 | 0.124239 | 0.23016 |
| COX5B | 1061 | 1121 | 631 | 903 | 528 | 630 | 409 | 409 | 857 | 1384 | 451 | 352 | 0.727224 | 0.40945 | 0.590437 | 0.300617 | -0.5281 |
| RPS17B | 2307 | 2659 | 1159 | 1809 | 2513 | 2020 | 4141 | 4492 | 4621 | 5765 | 5737 | 4407 | 0.364811 | 0.219681 | 0.309453 | 0.23743 | -0.49431 |
| RPS27B | 1964 | 2719 | 1359 | 2193 | 2391 | 1937 | 16178 | 15957 | 15716 | 23960 | 24736 | 24605 | 0.126923 | 0.081721 | 0.08056 | 0.655815 | 0.020653 |
| RPL27B | 2098 | 2648 | 1497 | 2016 | 2164 | 2386 | 8300 | 8720 | 7633 | 10549 | 10009 | 10097 | 0.217352 | 0.162205 | 0.184455 | 0.236763 | -0.18545 |
| RPL27A | 3333 | 3037 | 2196 | 3149 | 2675 | 2548 | 32790 | 33478 | 24217 | 31250 | 34002 | 34096 | 0.08772 | 0.087342 | 0.071234 | 0.300336 | 0.294113 |
| RPS27A | 3377 | 4145 | 2529 | 3496 | 4226 | 3003 | 1673 | 1383 | 1033 | 1231 | 1554 | 1447 | 0.709266 | 0.724788 | 0.702987 | 0.012829 | 0.044061 |
| NOG2 | 36807 | 34881 | 17084 | 40466 | 26939 | 32813 | 628 | 811 | 472 | 566 | 579 | 548 | 0.980251 | 0.97966 | 0.981266 | -0.00149 | -0.00236 |
| UBC5 | 135 | 102 | 93 | 201 | 95 | 108 | 467 | 380 | 710 | 743 | 575 | 375 | 0.217935 | 0.16437 | 0.182697 | 0.254449 | -0.15251 |
| DTD1 | 76 | 92 | 74 | 72 | 98 | 74 | 1073 | 1619 | 1525 | 2532 | 1907 | 1695 | 0.059957 | 0.036964 | 0.045355 | 0.402679 | -0.29512 |
| EPT1 | 201 | 137 | 189 | 248 | 242 | 176 | 609 | 688 | 616 | 815 | 687 | 653 | 0.207104 | 0.234042 | 0.2364 | -0.19087 | -0.01446 |
| YJR079W | 2493 | 2507 | 2366 | 2968 | 2032 | 1733 | 0 | 0 | 0 | 0 | 0 | 0 | 1 | 1 | 1 | 0 | 0 |
| YJR112W-A | 1584 | 1259 | 1039 | 1962 | 978 | 978 | 0 | 0 | 0 | 0 | 0 | 0 | 1 | 1 | 1 | 0 | 0 |
| NSP1 | 362 | 222 | 264 | 309 | 72 | 106 | 347 | 274 | 433 | 367 | 226 | 203 | 0.479079 | 0.417933 | 0.292326 | 0.712685 | 0.515693 |
| TEF4 | 4935 | 5006 | 1938 | 3104 | 2674 | 2739 | 47333 | 44246 | 25571 | 34239 | 36427 | 31970 | 0.098029 | 0.076786 | 0.07365 | 0.412519 | 0.060146 |
| TUB3 | 457 | 405 | 206 | 330 | 359 | 245 | 2988 | 2534 | 2242 | 2908 | 2098 | 1746 | 0.135229 | 0.093033 | 0.134583 | 0.006904 | -0.53269 |
| TUB1 | 53 | 67 | 77 | 101 | 84 | 76 | 1707 | 1365 | 1382 | 2054 | 1463 | 1266 | 0.038451 | 0.049822 | 0.055465 | -0.52858 | -0.15481 |
| YDR210W- | 27 | 25 | 33 | 63 | 13 | 8 | 414 | 398 | 352 | 435 | 58 | 152 | 0.060163 | 0.10611 | 0.116549 | -0.95399 | -0.13538 |

| | premRNA_ fast_A | premRNA_ fast_B | premRNA_ slow_A | premRNA_ slow_B | premRNA_ wt_A | premRNA_ wt_B | mRNA_ fast_A | mRNA_ fast_B | mRNA_ slow_A | mRNA_ slow_B | mRNA_ wt_A | mRNA_ wt_B | premRNA ratio_fast | premRNA ratio_slow | premRNA ratio_wt | premRNAra tio_fast/wt (log2) | premRNA ratio_slow/ wt(log2) |
|---|---|---|---|---|---|---|---|---|---|---|---|---|---|---|---|---|---|
| YOP1 | 644 | 724 | 316 | 711 | 842 | 650 | 3453 | 3176 | 3583 | 5283 | 4577 | 4152 | 0.171415 | 0.099833 | 0.14537 | 0.237763 | -0.54215 |
| YLL067C | 62 | 46 | 104 | 64 | 84 | 93 | 0 | 0 | 0 | 0 | 0 | 0 | 1 | 1 | 1 | 0 | 0 |
| SRC1 | 113 | 189 | 124 | 207 | 108 | 138 | 391 | 723 | 688 | 767 | 435 | 588 | 0.215722 | 0.182618 | 0.194489 | 0.149483 | -0.09086 |
| NMD2 | 91 | 107 | 101 | 116 | 35 | 75 | 236 | 430 | 325 | 454 | 87 | 87 | 0.238771 | 0.220299 | 0.374924 | -0.65097 | -0.76714 |
| AI2 | 43245 | 63639 | 232 | 352 | 51011 | 68457 | 260 | 359 | 1 | 2 | 408 | 506 | 0.994207 | 0.995029 | 0.992364 | 0.002677 | 0.00387 |
| YML6 | 97 | 105 | 66 | 71 | 48 | 88 | 417 | 325 | 314 | 561 | 384 | 217 | 0.216451 | 0.143013 | 0.199818 | 0.115355 | -0.48254 |
| NBL1 | 260 | 165 | 250 | 278 | 455 | 356 | 4270 | 4217 | 3412 | 3951 | 3404 | 2919 | 0.047525 | 0.067003 | 0.113304 | -1.25346 | -0.75791 |
| YDR210C-D | 12 | 31 | 91 | 197 | 69 | 45 | 259 | 359 | 900 | 1152 | 213 | 279 | 0.061884 | 0.11893 | 0.191785 | -1.63185 | -0.68937 |
| ECM33 | 1602 | 1580 | 1733 | 2277 | 1994 | 1986 | 4519 | 4135 | 4905 | 7384 | 6026 | 5518 | 0.269094 | 0.248381 | 0.256644 | 0.068342 | -0.04721 |
| YSF3 | 598 | 550 | 1112 | 1379 | 743 | 518 | 87 | 67 | 28 | 97 | 133 | 54 | 0.882201 | 0.95486 | 0.876884 | 0.008722 | 0.122904 |
| UBC12 | 357 | 431 | 652 | 867 | 430 | 371 | 120 | 148 | 157 | 211 | 117 | 109 | 0.746407 | 0.8051 | 0.779511 | -0.06261 | 0.046598 |
| UBC13 | 585 | 446 | 511 | 646 | 503 | 487 | 959 | 978 | 1425 | 1934 | 1137 | 1273 | 0.346044 | 0.257167 | 0.291706 | 0.246441 | -0.18181 |
| YLR202C | 247 | 337 | 218 | 255 | 273 | 298 | 0 | 0 | 0 | 0 | 0 | 0 | 1 | 1 | 1 | 0 | 0 |
| YOR318C | 1943 | 1865 | 1174 | 1146 | 1135 | 973 | 0 | 0 | 0 | 0 | 0 | 0 | 1 | 1 | 1 | 0 | 0 |
| MRK1 | 914 | 1078 | 291 | 549 | 597 | 508 | 7 | 36 | 5 | 31 | 117 | 28 | 0.980042 | 0.96483 | 0.891948 | 0.135884 | 0.113315 |
| RPL20B | 2770 | 3180 | 1395 | 2299 | 2435 | 2448 | 5246 | 5543 | 3568 | 4954 | 5648 | 5055 | 0.355056 | 0.299026 | 0.31376 | 0.178388 | -0.06939 |
| SPO22 | 97 | 110 | 107 | 133 | 76 | 154 | 3 | 0 | 0 | 0 | 0 | 0 | 0.985 | 1 | 1 | -0.0218 | 0 |
| RPL20A | 2508 | 3058 | 1629 | 2222 | 2077 | 2650 | 6677 | 7391 | 5268 | 7502 | 8265 | 6804 | 0.282857 | 0.232348 | 0.240568 | 0.233626 | -0.05016 |
| RPS16B | 2875 | 2873 | 1517 | 2319 | 2851 | 2554 | 24775 | 24828 | 18867 | 24463 | 29972 | 26523 | 0.103846 | 0.080505 | 0.087348 | 0.249609 | -0.1177 |
| RPL17A | 2039 | 2081 | 1257 | 1780 | 2187 | 1937 | 38087 | 34432 | 25028 | 30360 | 32567 | 28388 | 0.053904 | 0.051602 | 0.063401 | -0.23412 | -0.29708 |
| RPL17B | 874 | 852 | 885 | 1411 | 1581 | 1687 | 19269 | 19379 | 19905 | 23563 | 23309 | 20512 | 0.042752 | 0.049534 | 0.069757 | -0.70636 | -0.49393 |
| RPS16A | 2909 | 3290 | 2137 | 3259 | 2995 | 3108 | 9436 | 10175 | 9242 | 13327 | 13092 | 12440 | 0.23999 | 0.192147 | 0.193036 | 0.314101 | -0.00666 |
| RPL26A | 2567 | 2830 | 1823 | 2712 | 3471 | 2603 | 16077 | 15053 | 15999 | 21242 | 23946 | 20881 | 0.147968 | 0.107753 | 0.118721 | 0.317711 | -0.13984 |
| RPL26B | 2837 | 3932 | 2543 | 3538 | 2349 | 2725 | 32691 | 32373 | 29412 | 38293 | 41235 | 40092 | 0.094079 | 0.08208 | 0.058769 | 0.6788 | 0.481957 |
| SEC14 | 662 | 529 | 582 | 781 | 577 | 731 | 1747 | 1327 | 1618 | 2032 | 1453 | 1565 | 0.279912 | 0.271092 | 0.301308 | -0.10627 | -0.15245 |
| RPS7A | 3545 | 3785 | 2635 | 3514 | 3450 | 3321 | 77457 | 76820 | 48960 | 68065 | 74992 | 65096 | 0.045361 | 0.050082 | 0.046261 | -0.02835 | 0.114486 |
| SEC17 | 244 | 323 | 367 | 330 | 372 | 288 | 915 | 1256 | 1106 | 1591 | 823 | 805 | 0.207543 | 0.210468 | 0.287396 | -0.46963 | -0.44944 |
| GIM5 | 116 | 186 | 235 | 335 | 194 | 172 | 592 | 563 | 716 | 817 | 658 | 569 | 0.206086 | 0.268953 | 0.229909 | -0.15781 | 0.226293 |
| TAN1 | 134 | 193 | 323 | 332 | 300 | 234 | 267 | 339 | 969 | 1178 | 816 | 814 | 0.348473 | 0.234934 | 0.24605 | 0.502098 | -0.0667 |
| VPS75 | 358 | 403 | 204 | 430 | 396 | 427 | 1186 | 1551 | 905 | 1265 | 987 | 941 | 0.219054 | 0.218818 | 0.299234 | -0.44999 | -0.45154 |
| RPL6A | 1869 | 1736 | 941 | 1284 | 1294 | 1219 | 10228 | 9716 | 8809 | 12512 | 13318 | 11724 | 0.153045 | 0.094792 | 0.09137 | 0.744169 | 0.053043 |
| RPS7B | 2022 | 1952 | 1568 | 2366 | 3173 | 2610 | 27095 | 23675 | 21254 | 22776 | 30048 | 24798 | 0.072807 | 0.081406 | 0.09537 | -0.38946 | -0.2284 |
| YPR063C | 129 | 136 | 60 | 107 | 135 | 107 | 1347 | 1385 | 1243 | 1853 | 1349 | 1086 | 0.088407 | 0.05032 | 0.09033 | -0.03105 | -0.84408 |
| BET4 | 140 | 63 | 104 | 188 | 193 | 173 | 86 | 149 | 59 | 90 | 54 | 102 | 0.458319 | 0.657148 | 0.705234 | -0.62175 | -0.10188 |
| HAC1 | 9598 | 9551 | 8447 | 12580 | 12406 | 10877 | 0 | 0 | 2 | 3 | 20 | 0 | 1 | 0.999762 | 0.999195 | 0.001161 | 0.000819 |
| SRB2 | 257 | 298 | 283 | 353 | 362 | 303 | 1528 | 1171 | 978 | 1191 | 1336 | 883 | 0.173418 | 0.226526 | 0.234336 | -0.43432 | -0.0489 |
| YGR109W- | 0 | 7 | 2 | 5 | 3 | 7 | 162 | 297 | 285 | 397 | 367 | 299 | 0.011513 | 0.009703 | 0.015492 | -0.42824 | -0.67498 |
| POP8 | 105 | 160 | 131 | 191 | 104 | 105 | 486 | 610 | 930 | 1257 | 840 | 923 | 0.192729 | 0.127687 | 0.106155 | 0.860401 | 0.266445 |
| HNT2 | 319 | 344 | 134 | 323 | 243 | 319 | 457 | 454 | 454 | 570 | 365 | 373 | 0.42108 | 0.294797 | 0.430327 | -0.03134 | -0.54571 |
| ERV1 | 608 | 812 | 633 | 910 | 947 | 716 | 1946 | 1714 | 1423 | 1931 | 1702 | 1477 | 0.279757 | 0.314095 | 0.341993 | -0.28979 | -0.12277 |
| HNT1 | 856 | 869 | 993 | 1201 | 1045 | 1102 | 12555 | 11518 | 7318 | 10407 | 9005 | 7432 | 0.066991 | 0.111472 | 0.116555 | -0.79897 | -0.06434 |

| | premRNA_ fast_A | premRNA_ fast_B | premRNA_ slow_A | premRNA_ slow_B | premRNA_ wt_A | premRNA_ wt_B | mRNA_ fast_A | mRNA_ fast_B | mRNA_ slow_A | mRNA_ slow_B | mRNA_ wt_A | mRNA_ wt_B | premRNA ratio_fast | premRNA ratio_slow | premRNA ratio_wt | premRNAra tio_fast/wt (log2) | premRNA ratio_slow/ wt(log2) |
|---|---|---|---|---|---|---|---|---|---|---|---|---|---|---|---|---|---|
| YNL050C | 105 | 192 | 187 | 216 | 108 | 163 | 78 | 123 | 182 | 189 | 139 | 132 | 0.591647 | 0.520054 | 0.494895 | 0.257616 | 0.07154 |
| YBL005W-E | 70 | 38 | 63 | 214 | 43 | 62 | 1415 | 782 | 3141 | 4265 | 1241 | 939 | 0.04674 | 0.033721 | 0.047714 | -0.02975 | -0.50076 |
| RPS14A | 5304 | 6061 | 1692 | 2339 | 2619 | 2663 | 31736 | 26432 | 22857 | 27777 | 29542 | 27726 | 0.164865 | 0.073295 | 0.084532 | 0.963708 | -0.20579 |
| YBR062C | 248 | 280 | 365 | 530 | 236 | 320 | 409 | 598 | 390 | 658 | 309 | 322 | 0.34819 | 0.464786 | 0.465735 | -0.41963 | -0.00294 |
| YML133C | 577 | 625 | 417 | 596 | 433 | 613 | 0 | 0 | 0 | 0 | 0 | 0 | 1 | 1 | 1 | 0 | 0 |
| YDR381C-A | 323 | 289 | 442 | 567 | 280 | 285 | 334 | 208 | 283 | 225 | 86 | 147 | 0.536559 | 0.662782 | 0.712375 | -0.4089 | -0.1041 |
| RPL18A | 2560 | 2926 | 1466 | 2369 | 2847 | 2976 | 55533 | 55818 | 33450 | 49158 | 53738 | 48442 | 0.046938 | 0.043981 | 0.054096 | -0.20476 | -0.29864 |
| RPL35A | 3773 | 3798 | 3452 | 4739 | 4943 | 4026 | 6139 | 5734 | 6292 | 7094 | 8010 | 6430 | 0.389549 | 0.37738 | 0.383326 | 0.02323 | -0.02256 |
| RPL35B | 3061 | 3662 | 2386 | 3629 | 3131 | 2753 | 7343 | 7159 | 5349 | 6672 | 7275 | 7138 | 0.316315 | 0.330382 | 0.289609 | 0.127255 | 0.190029 |
| RPL18B | 2398 | 2636 | 2222 | 3114 | 3300 | 3152 | 5431 | 5147 | 3943 | 5682 | 6333 | 6272 | 0.322492 | 0.357223 | 0.338519 | -0.06997 | 0.07759 |
| DBP2 | 823 | 911 | 454 | 820 | 748 | 773 | 2266 | 2390 | 879 | 1297 | 1721 | 1383 | 0.271203 | 0.363963 | 0.330745 | -0.28635 | 0.13807 |
| YPR202W | 889 | 997 | 780 | 1287 | 1000 | 890 | 43 | 52 | 27 | 42 | 12 | 23 | 0.952146 | 0.96747 | 0.981475 | -0.04377 | -0.02074 |
| YHL009W-I | 0 | 0 | 0 | 10 | 0 | 0 | 126 | 110 | 121 | 191 | 51 | 43 | 0 | 0.024876 | 0 | | inf |
| TAF14 | 638 | 706 | 1019 | 1159 | 760 | 735 | 433 | 485 | 361 | 352 | 277 | 356 | 0.594242 | 0.752724 | 0.703289 | -0.24307 | 0.098004 |
| PFY1 | 1349 | 1259 | 1168 | 1834 | 1835 | 1527 | 6920 | 6062 | 6727 | 8500 | 8193 | 7303 | 0.167555 | 0.162707 | 0.17796 | -0.08692 | -0.12928 |
| SAR1 | 218 | 180 | 299 | 355 | 237 | 243 | 4008 | 3755 | 4494 | 5745 | 4240 | 3367 | 0.048664 | 0.06029 | 0.060125 | -0.3051 | 0.003943 |
| DCN1 | 61 | 65 | 146 | 113 | 71 | 89 | 24 | 11 | 27 | 53 | 54 | 31 | 0.786455 | 0.762327 | 0.654833 | 0.264237 | 0.219282 |
| PSP2 | 211 | 276 | 215 | 268 | 170 | 272 | 165 | 220 | 182 | 223 | 112 | 115 | 0.558811 | 0.543693 | 0.65284 | -0.22437 | -0.26394 |
| PRE3 | 573 | 480 | 444 | 661 | 1159 | 1084 | 4173 | 3881 | 3669 | 5165 | 3687 | 3027 | 0.1154 | 0.110704 | 0.251425 | -1.12348 | -1.18342 |
| MPT5 | 1251 | 1171 | 907 | 979 | 756 | 705 | 14 | 0 | 29 | 40 | 33 | 10 | 0.994466 | 0.964881 | 0.972094 | 0.032826 | -0.01074 |
| ECM9 | 55 | 162 | 122 | 218 | 122 | 138 | 323 | 505 | 793 | 941 | 575 | 569 | 0.194191 | 0.160713 | 0.185113 | 0.069064 | -0.20392 |
| YLR227W-E | 69 | 43 | 280 | 276 | 84 | 101 | 74 | 63 | 205 | 331 | 101 | 94 | 0.444089 | 0.516007 | 0.486001 | -0.13011 | 0.086431 |
| YPR170W-I | 319 | 445 | 215 | 267 | 174 | 245 | 10897 | 10770 | 7646 | 10082 | 9824 | 8662 | 0.03406 | 0.026575 | 0.022455 | 0.601055 | 0.24303 |
| MCM21 | 109 | 115 | 163 | 178 | 102 | 106 | 253 | 357 | 350 | 443 | 326 | 252 | 0.272375 | 0.302187 | 0.267204 | 0.027652 | 0.177501 |
| YCL002C | 525 | 500 | 563 | 782 | 981 | 788 | 216 | 353 | 114 | 161 | 209 | 120 | 0.647334 | 0.830439 | 0.846106 | -0.38633 | -0.02696 |
| TFC3 | 46 | 117 | 50 | 181 | 179 | 171 | 388 | 551 | 628 | 701 | 410 | 369 | 0.14057 | 0.139481 | 0.310286 | -1.14231 | -1.15353 |
| RPL13A | 1464 | 1385 | 1182 | 1479 | 1750 | 1545 | 3803 | 4081 | 3445 | 4377 | 4717 | 3886 | 0.265671 | 0.254009 | 0.277541 | -0.06306 | -0.12782 |
| RPL13B | 3017 | 3210 | 1613 | 2140 | 2257 | 2107 | 5910 | 6838 | 4582 | 7116 | 8128 | 6312 | 0.328715 | 0.245786 | 0.2338 | 0.491563 | 0.07213 |
| RUB1 | 568 | 537 | 583 | 853 | 678 | 657 | 3787 | 4262 | 4413 | 4678 | 2846 | 3312 | 0.121162 | 0.135458 | 0.178964 | -0.56274 | -0.40183 |
| YNL284C-B | 0 | 1 | 6 | 21 | 1 | 6 | 287 | 427 | 1639 | 1886 | 285 | 230 | 0.001168 | 0.00733 | 0.01446 | -3.62969 | -0.98025 |
| RPL40A | 11578 | 11571 | 9516 | 12981 | 15202 | 11219 | 12834 | 13138 | 13110 | 17679 | 19349 | 15927 | 0.471283 | 0.421982 | 0.426635 | 0.143589 | -0.01582 |
| SCS22 | 273 | 139 | 322 | 459 | 341 | 212 | 214 | 407 | 480 | 515 | 220 | 218 | 0.407577 | 0.436374 | 0.550433 | -0.4335 | -0.335 |
| RPL22A | 6370 | 7535 | 3731 | 6295 | 6333 | 6789 | 11845 | 12027 | 11417 | 13655 | 17459 | 16174 | 0.367449 | 0.280921 | 0.280916 | 0.387405 | 2.71E-05 |
| RPL22B | 11564 | 13487 | 7976 | 12443 | 13820 | 12781 | 384 | 309 | 532 | 649 | 697 | 683 | 0.972731 | 0.943949 | 0.95063 | 0.033158 | -0.01017 |
| QCR9 | 390 | 452 | 226 | 381 | 606 | 493 | 741 | 894 | 693 | 1160 | 634 | 742 | 0.340319 | 0.246581 | 0.44395 | -0.38351 | -0.84834 |
| YHL050C | 1013 | 880 | 919 | 1744 | 1120 | 1151 | 0 | 0 | 0 | 0 | 0 | 0 | 1 | 1 | 1 | 0 | 0 |
| RPL40B | 1335 | 1314 | 810 | 1473 | 1725 | 1690 | 16250 | 16269 | 15322 | 19020 | 22130 | 19643 | 0.075324 | 0.061044 | 0.075766 | -0.00844 | -0.31169 |
| BUD25 | 1712 | 1614 | 1158 | 1535 | 1032 | 971 | 0 | 0 | 0 | 0 | 0 | 0 | 1 | 1 | 1 | 0 | 0 |
| RPL36A | 3231 | 4022 | 2693 | 3789 | 3071 | 3282 | 8508 | 8131 | 9153 | 12329 | 10654 | 10768 | 0.303092 | 0.231206 | 0.228673 | 0.406467 | 0.015894 |
| RPS4A | 1222 | 1514 | 825 | 1253 | 1516 | 1494 | 24674 | 22558 | 21900 | 26360 | 29382 | 23970 | 0.055042 | 0.04084 | 0.053868 | 0.0311 | -0.39943 |
| RPS14B | 3614 | 3997 | 2598 | 3947 | 6078 | 5713 | 3418 | 2511 | 1744 | 1988 | 4471 | 3473 | 0.564052 | 0.63169 | 0.599047 | -0.08684 | 0.076548 |

| | premRNA_fast_A | premRNA_fast_B | premRNA_slow_A | premRNA_slow_B | premRNA_wt_A | premRNA_wt_B | mRNA_fast_A | mRNA_fast_B | mRNA_slow_A | mRNA_slow_B | mRNA_wt_A | mRNA_wt_B | premRNA ratio_fast | premRNA ratio_slow | premRNA ratio_wt | premRNAratio_fast/wt (log2) | premRNA ratio_slow/wt(log2) |
|---|---|---|---|---|---|---|---|---|---|---|---|---|---|---|---|---|---|
| RPL36B | 1373 | 1313 | 894 | 961 | 1365 | 1132 | 34145 | 31134 | 29370 | 36932 | 43261 | 35144 | 0.039561 | 0.02745 | 0.030896 | 0.356649 | -0.17061 |
| RPS30B | 1416 | 1744 | 1087 | 2059 | 1452 | 1471 | 7751 | 7140 | 5288 | 8186 | 8001 | 7222 | 0.175388 | 0.185743 | 0.161409 | 0.119822 | 0.202584 |
| RPS30A | 3479 | 4608 | 1919 | 3777 | 2680 | 2778 | 23433 | 22750 | 17665 | 21855 | 21698 | 15351 | 0.148853 | 0.122672 | 0.131585 | 0.177894 | -0.1012 |
| YFR045W | 117 | 48 | 105 | 164 | 67 | 47 | 83 | 97 | 120 | 146 | 69 | 56 | 0.458017 | 0.497849 | 0.474479 | -0.05094 | 0.069366 |
| BIG1 | 148 | 83 | 115 | 126 | 140 | 95 | 552 | 736 | 569 | 768 | 391 | 537 | 0.156386 | 0.154534 | 0.206985 | -0.40442 | -0.4216 |
| RPL19B | 7766 | 7450 | 7081 | 9380 | 9475 | 9517 | 6090 | 5295 | 5504 | 6763 | 6282 | 6120 | 0.572511 | 0.571855 | 0.60497 | -0.07956 | -0.08121 |
| SFT1 | 431 | 437 | 343 | 499 | 440 | 432 | 445 | 562 | 714 | 839 | 467 | 502 | 0.464723 | 0.348724 | 0.473821 | -0.02797 | -0.44226 |
| RPL19A | 2941 | 3235 | 2065 | 2987 | 3608 | 3476 | 2903 | 2940 | 2668 | 3606 | 3721 | 2728 | 0.513569 | 0.444677 | 0.526287 | -0.03529 | -0.24309 |
| QCR10 | 37 | 99 | 85 | 104 | 45 | 64 | 213 | 264 | 227 | 293 | 198 | 164 | 0.210364 | 0.2672 | 0.232943 | -0.14709 | 0.197942 |
| DID4 | 31 | 162 | 151 | 158 | 115 | 125 | 490 | 468 | 598 | 676 | 433 | 403 | 0.158322 | 0.195525 | 0.223298 | -0.49611 | -0.19162 |
| RPL6B | 1880 | 2029 | 1415 | 1676 | 2143 | 1619 | 18820 | 19719 | 15785 | 20311 | 26280 | 21313 | 0.092059 | 0.079247 | 0.072998 | 0.334688 | 0.118495 |
| YOL047C | 183 | 174 | 344 | 454 | 179 | 164 | 0 | 7 | 0 | 6 | 39 | 3 | 0.980663 | 0.993478 | 0.901568 | 0.12132 | 0.140051 |
| YOL103W-I | 0 | 14 | 2 | 2 | 10 | 4 | 40 | 35 | 68 | 118 | 38 | 91 | 0.142857 | 0.022619 | 0.125219 | 0.190116 | -2.46885 |
| MOB2 | 47 | 64 | 80 | 104 | 86 | 95 | 69 | 34 | 31 | 65 | 59 | 33 | 0.529117 | 0.668053 | 0.667645 | -0.3355 | 0.00088 |
| KEI1 | 148 | 167 | 165 | 240 | 171 | 148 | 548 | 461 | 597 | 637 | 562 | 396 | 0.239284 | 0.245098 | 0.252673 | -0.07855 | -0.04392 |
| CMC2 | 65 | 84 | 99 | 142 | 97 | 138 | 482 | 622 | 1442 | 1659 | 552 | 643 | 0.118905 | 0.071545 | 0.163079 | -0.45576 | -1.18865 |
| CMC4 | 176 | 127 | 217 | 222 | 220 | 129 | 76 | 82 | 175 | 183 | 134 | 87 | 0.653034 | 0.55086 | 0.609346 | 0.099898 | -0.14558 |
| YIL082W-A | 2 | 11 | 8 | 5 | 4 | 13 | 559 | 532 | 275 | 659 | 418 | 346 | 0.011911 | 0.017899 | 0.022845 | -0.93954 | -0.35198 |
| IMD4 | 3148 | 3598 | 1521 | 2360 | 2349 | 2220 | 8303 | 9440 | 5532 | 8894 | 7927 | 7556 | 0.275437 | 0.212678 | 0.227839 | 0.273706 | -0.09934 |
| GOT1 | 193 | 187 | 167 | 249 | 186 | 161 | 1119 | 1077 | 997 | 1651 | 1333 | 1385 | 0.147523 | 0.137262 | 0.113294 | 0.380867 | 0.276853 |
| STO1 | 507 | 579 | 665 | 601 | 593 | 444 | 588 | 669 | 490 | 645 | 484 | 335 | 0.463478 | 0.529051 | 0.560283 | -0.27365 | -0.08275 |
| ERV41 | 162 | 182 | 150 | 222 | 213 | 214 | 1303 | 1122 | 1394 | 1424 | 1309 | 1082 | 0.125075 | 0.116011 | 0.152535 | -0.28635 | -0.39488 |
| YPR153W | 163 | 172 | 452 | 462 | 310 | 297 | 388 | 500 | 401 | 316 | 430 | 281 | 0.275889 | 0.561862 | 0.46638 | -0.75742 | 0.268711 |
| YCL019W | 2 | 0 | 0 | 3 | 0 | 0 | 28 | 113 | 167 | 413 | 60 | 63 | 0.033333 | 0.003606 | 0 | inf | inf |
| YLL066C | 76 | 65 | 76 | 56 | 54 | 64 | 2 | 0 | 0 | 0 | 0 | 0 | 0.987179 | 1 | 1 | -0.01862 | 0 |
| CNB1 | 239 | 152 | 232 | 394 | 126 | 127 | 1548 | 1626 | 1949 | 2571 | 1525 | 1511 | 0.109617 | 0.119628 | 0.076925 | 0.510932 | 0.637027 |
| OSW2 | 82 | 81 | 66 | 77 | 63 | 97 | 0 | 0 | 0 | 0 | 0 | 0 | 1 | 1 | 1 | 0 | 0 |
| SAC6 | 85 | 150 | 148 | 153 | 83 | 108 | 2022 | 1702 | 1955 | 2719 | 2056 | 1723 | 0.060668 | 0.061824 | 0.048894 | 0.311279 | 0.338527 |
| SNC1 | 232 | 97 | 142 | 286 | 198 | 280 | 3344 | 2785 | 2578 | 3437 | 3106 | 2649 | 0.049267 | 0.064513 | 0.077762 | -0.65843 | -0.26947 |
| EST3 | 43 | 33 | 18 | 17 | 70 | 35 | 1914 | 1763 | 1544 | 2137 | 2066 | 1891 | 0.020173 | 0.009708 | 0.025472 | -0.33646 | -1.39167 |
| GLC7 | 584 | 498 | 400 | 692 | 272 | 456 | 1721 | 1912 | 3037 | 4242 | 1838 | 2254 | 0.230001 | 0.128316 | 0.148588 | 0.630322 | -0.21162 |
| YDL012C | 385 | 414 | 523 | 490 | 409 | 421 | 1846 | 1649 | 1777 | 2001 | 1438 | 968 | 0.186623 | 0.21205 | 0.262268 | -0.49091 | -0.30664 |
| GIM4 | 256 | 290 | 396 | 873 | 242 | 276 | 977 | 881 | 836 | 1525 | 1057 | 944 | 0.227638 | 0.342741 | 0.206263 | 0.142252 | 0.732631 |
| RPL21A | 9477 | 9451 | 4750 | 6720 | 7641 | 7104 | 19766 | 19305 | 14565 | 19650 | 22326 | 21020 | 0.32637 | 0.250379 | 0.253788 | 0.362883 | -0.01951 |
| YBR219C | 961 | 1028 | 729 | 1039 | 1165 | 882 | 0 | 0 | 0 | 0 | 0 | 0 | 1 | 1 | 1 | 0 | 0 |
| SMD2 | 27 | 84 | 59 | 38 | 73 | 71 | 52 | 83 | 161 | 116 | 83 | 47 | 0.422383 | 0.257468 | 0.534822 | -0.34051 | -1.05467 |
| RPL21B | 6573 | 6264 | 3360 | 5560 | 6075 | 5277 | 14176 | 12659 | 9617 | 13305 | 14382 | 13667 | 0.323906 | 0.276823 | 0.287761 | 0.170704 | -0.05591 |
| RPS13 | 32757 | 33391 | 13861 | 20174 | 25585 | 21921 | 12713 | 12960 | 10449 | 14866 | 17348 | 14839 | 0.720402 | 0.572959 | 0.596128 | 0.273179 | -0.05719 |
| RPP1B | 3459 | 3348 | 2721 | 4154 | 4685 | 5164 | 61825 | 60268 | 45962 | 65371 | 62118 | 59455 | 0.052806 | 0.05782 | 0.075023 | -0.50663 | -0.37576 |
| MUD1 | 31 | 67 | 45 | 89 | 65 | 60 | 46 | 72 | 81 | 96 | 36 | 52 | 0.442306 | 0.419112 | 0.589639 | -0.41479 | -0.4925 |
| RPS23B | 2643 | 2903 | 1728 | 2786 | 2817 | 2721 | 47937 | 48544 | 34209 | 46954 | 46921 | 45832 | 0.05434 | 0.052048 | 0.056339 | -0.05212 | -0.11431 |

| | premRNA_ fast_A | premRNA_ fast_B | premRNA_ slow_A | premRNA_ slow_B | premRNA_ wt_A | premRNA_ wt_B | mRNA_ fast_A | mRNA_ fast_B | mRNA_ slow_A | mRNA_ slow_B | mRNA_ wt_A | mRNA_ wt_B | premRNA ratio_fast | premRNA ratio_slow | premRNA ratio_wt | premRNAratio_fast/wt (log2) | premRNA ratio_slow/ wt(log2) |
|---|---|---|---|---|---|---|---|---|---|---|---|---|---|---|---|---|---|
| RPL23B | 6219 | 6117 | 3896 | 5868 | 6985 | 6069 | 38703 | 35364 | 33114 | 42844 | 49086 | 40374 | 0.142953 | 0.112866 | 0.127625 | 0.163622 | -0.1773 |
| RPL23A | 2696 | 2928 | 2252 | 2403 | 2900 | 2658 | 24928 | 26625 | 23757 | 33011 | 29283 | 28051 | 0.098336 | 0.07722 | 0.088332 | 0.154786 | -0.19396 |
| RPS23A | 2079 | 2254 | 1471 | 2387 | 2630 | 2454 | 23914 | 21271 | 18446 | 26058 | 23788 | 21631 | 0.087898 | 0.078886 | 0.100721 | -0.19647 | -0.35252 |
| RPS9B | 8394 | 9030 | 4237 | 5585 | 9919 | 8361 | 25405 | 24513 | 18985 | 22399 | 27688 | 24705 | 0.258779 | 0.191017 | 0.258306 | 0.002637 | -0.43538 |
| RPS9A | 4568 | 4763 | 3262 | 4822 | 5716 | 6217 | 212 | 219 | 839 | 975 | 996 | 849 | 0.955845 | 0.813613 | 0.865728 | 0.142863 | -0.08957 |
| PMI40 | 137 | 98 | 79 | 143 | 117 | 128 | 2069 | 2112 | 1664 | 2429 | 2254 | 2086 | 0.053224 | 0.050461 | 0.05358 | -0.00963 | -0.08652 |
| RPS4B | 1363 | 1384 | 561 | 1058 | 1308 | 996 | 13148 | 13605 | 8297 | 13005 | 12833 | 11929 | 0.093132 | 0.069283 | 0.084778 | 0.135572 | -0.2912 |
| OM14 | 67 | 131 | 105 | 191 | 151 | 218 | 309 | 266 | 346 | 481 | 386 | 290 | 0.254083 | 0.258521 | 0.355163 | -0.48318 | -0.4582 |
| RRT8 | 57 | 133 | 25 | 84 | 89 | 47 | 311 | 382 | 362 | 487 | 318 | 292 | 0.206572 | 0.105855 | 0.178658 | 0.209442 | -0.75511 |
| PHO85 | 186 | 100 | 154 | 171 | 209 | 181 | 249 | 234 | 553 | 408 | 158 | 192 | 0.363494 | 0.256579 | 0.527368 | -0.53688 | -1.03941 |
| YHR097C | 102 | 91 | 56 | 156 | 64 | 65 | 314 | 217 | 277 | 385 | 201 | 235 | 0.270323 | 0.228262 | 0.229088 | 0.238784 | -0.00521 |
| YSC84 | 157 | 184 | 110 | 249 | 187 | 227 | 587 | 654 | 443 | 723 | 278 | 368 | 0.215296 | 0.227544 | 0.391832 | -0.86391 | -0.78409 |
| GCR1 | 2773 | 3145 | 4598 | 6509 | 3655 | 3973 | 0 | 0 | 0 | 0 | 0 | 0 | 1 | 1 | 1 | 0 | 0 |
| YCL012C | 45 | 22 | 34 | 25 | 9 | 73 | 612 | 599 | 527 | 561 | 485 | 457 | 0.05196 | 0.051634 | 0.077977 | -0.58565 | -0.59473 |
| RPL28 | 74219 | 81440 | 37112 | 60086 | 61627 | 58861 | 64685 | 64190 | 53985 | 76206 | 72229 | 72968 | 0.546772 | 0.424126 | 0.453446 | 0.270007 | -0.09644 |
| RPL25 | 7090 | 7022 | 6273 | 8017 | 9714 | 9022 | 16401 | 15748 | 13049 | 16963 | 18551 | 17237 | 0.305103 | 0.322796 | 0.343627 | -0.17155 | -0.09022 |
| SPO1 | 447 | 505 | 339 | 469 | 309 | 335 | 25 | 18 | 0 | 37 | 7 | 0 | 0.956309 | 0.963439 | 0.988924 | -0.04838 | -0.03767 |
| VMA10 | 371 | 301 | 586 | 706 | 490 | 435 | 850 | 1084 | 1270 | 1364 | 1256 | 841 | 0.260589 | 0.328398 | 0.310775 | -0.2541 | 0.079573 |
| VPS29 | 56 | 123 | 113 | 269 | 131 | 176 | 773 | 855 | 954 | 1673 | 1045 | 1110 | 0.096659 | 0.122211 | 0.124127 | -0.36083 | -0.02244 |
| YPR098C | 100 | 243 | 140 | 247 | 65 | 119 | 1095 | 1209 | 1096 | 1592 | 941 | 926 | 0.125519 | 0.12379 | 0.089244 | 0.492076 | 0.472073 |
| MMS2 | 84 | 74 | 80 | 145 | 99 | 69 | 216 | 384 | 980 | 1230 | 564 | 628 | 0.220786 | 0.090463 | 0.124158 | 0.830466 | -0.45678 |
| ABP140 | 5 | 25 | 3 | 22 | 26 | 7 | 542 | 786 | 832 | 1081 | 678 | 764 | 0.019983 | 0.011769 | 0.023005 | -0.20317 | -0.96696 |
| YEL076C-A | 603 | 509 | 519 | 747 | 478 | 550 | 0 | 0 | 0 | 0 | 0 | 0 | 1 | 1 | 1 | 0 | 0 |
| YIP3 | 189 | 311 | 230 | 235 | 175 | 208 | 801 | 723 | 748 | 848 | 840 | 760 | 0.245841 | 0.226082 | 0.193645 | 0.344314 | 0.223431 |
| YPL109C | 124 | 245 | 122 | 285 | 123 | 207 | 182 | 245 | 306 | 314 | 268 | 148 | 0.452614 | 0.38042 | 0.448838 | 0.012087 | -0.2386 |
| OST5 | 181 | 156 | 207 | 295 | 264 | 261 | 1970 | 1642 | 1908 | 2677 | 2354 | 1990 | 0.085455 | 0.098566 | 0.108394 | -0.34305 | -0.13713 |
| RPS0A | 2248 | 2230 | 1165 | 1730 | 2037 | 1714 | 20993 | 18887 | 17594 | 23039 | 23527 | 20927 | 0.101164 | 0.065974 | 0.077693 | 0.38084 | -0.23587 |
| RPS0B | 3436 | 3944 | 2107 | 2795 | 4120 | 2971 | 28071 | 25287 | 17741 | 25125 | 24649 | 24511 | 0.12199 | 0.103132 | 0.125658 | -0.04274 | -0.28501 |
| APS3 | 109 | 142 | 132 | 379 | 174 | 326 | 464 | 425 | 557 | 686 | 408 | 481 | 0.220334 | 0.273725 | 0.351467 | -0.6737 | -0.36066 |
| YDR261C-D | 11 | 42 | 102 | 187 | 41 | 40 | 44 | 67 | 186 | 198 | 37 | 66 | 0.292661 | 0.41994 | 0.4515 | -0.6255 | -0.10454 |
| YJL113W | 0 | 1 | 0 | 0 | 0 | 0 | 62 | 28 | 171 | 91 | 61 | 30 | 0.017241 | 0 | 0 | inf | |
| NCE101 | 714 | 629 | 689 | 812 | 454 | 391 | 797 | 768 | 1007 | 1386 | 804 | 731 | 0.461393 | 0.387838 | 0.354688 | 0.379446 | 0.128907 |
| PCH2 | 1323 | 1359 | 753 | 1127 | 865 | 947 | 10 | 2 | 6 | 24 | 9 | 12 | 0.995514 | 0.985622 | 0.988595 | 0.010063 | -0.00435 |
| YPR010C-A | 55 | 93 | 105 | 111 | 152 | 77 | 894 | 835 | 595 | 824 | 639 | 461 | 0.079086 | 0.134358 | 0.167642 | -1.0839 | -0.3193 |
| RPL34B | 2486 | 2989 | 1943 | 3279 | 2795 | 2566 | 18937 | 18047 | 20883 | 26308 | 27757 | 22643 | 0.129067 | 0.097974 | 0.096636 | 0.41748 | 0.019835 |
| RPL34A | 3608 | 3709 | 2601 | 3807 | 3104 | 3089 | 33161 | 25903 | 27787 | 30446 | 37319 | 30105 | 0.11169 | 0.098368 | 0.084923 | 0.395261 | 0.21203 |
| LSM7 | 149 | 129 | 121 | 329 | 194 | 178 | 719 | 870 | 951 | 1109 | 652 | 823 | 0.150394 | 0.170832 | 0.203568 | -0.43677 | -0.25294 |
| RPL43A | 8885 | 8716 | 5671 | 7026 | 8053 | 7478 | 14611 | 14106 | 10330 | 13222 | 15620 | 12328 | 0.380031 | 0.350706 | 0.358869 | 0.082657 | -0.0332 |
| LSM2 | 543 | 584 | 455 | 798 | 564 | 476 | 1625 | 1947 | 1791 | 3046 | 2200 | 2406 | 0.2406 | 0.205089 | 0.184608 | 0.382175 | 0.15179 |
| RPL43B | 3053 | 3135 | 3584 | 4954 | 4634 | 3965 | 2577 | 2367 | 1921 | 2675 | 2807 | 2542 | 0.556033 | 0.650204 | 0.616055 | -0.14789 | 0.077835 |
| RPS24B | 1499 | 1589 | 885 | 1749 | 2196 | 1859 | 4255 | 3174 | 2362 | 3619 | 3516 | 3342 | 0.297064 | 0.299189 | 0.370943 | -0.32042 | -0.31014 |

| | premRNA_fast_A | premRNA_fast_B | premRNA_slow_A | premRNA_slow_B | premRNA_wt_A | premRNA_wt_B | mRNA_fast_A | mRNA_fast_B | mRNA_slow_A | mRNA_slow_B | mRNA_wt_A | mRNA_wt_B | premRNA ratio_fast | premRNA ratio_slow | premRNA ratio_wt | premRNAratio_fast/wt (log2) | premRNA ratio_slow/wt(log2) |
|---|---|---|---|---|---|---|---|---|---|---|---|---|---|---|---|---|---|
| RPS24A | 6335 | 7118 | 3099 | 4826 | 4982 | 4635 | 10687 | 10767 | 8193 | 10148 | 10496 | 9874 | 0.385076 | 0.298367 | 0.320667 | 0.26407 | -0.10399 |
| REC102 | 243 | 443 | 106 | 216 | 230 | 229 | 16 | 0 | 12 | 9 | 19 | 9 | 0.969112 | 0.929153 | 0.94294 | 0.039498 | -0.02125 |
| PCC1 | 107 | 105 | 81 | 245 | 175 | 119 | 1268 | 1133 | 1243 | 1540 | 1077 | 854 | 0.081316 | 0.099217 | 0.131039 | -0.68838 | -0.40135 |
| REC107 | 1398 | 1669 | 947 | 1591 | 1331 | 1312 | 118 | 124 | 127 | 124 | 21 | 125 | 0.926503 | 0.904724 | 0.94874 | -0.03422 | -0.06854 |
| NYV1 | 132 | 132 | 103 | 227 | 210 | 218 | 551 | 433 | 653 | 610 | 355 | 439 | 0.213447 | 0.203725 | 0.351746 | -0.72066 | -0.78791 |
| CGI121 | 321 | 397 | 280 | 493 | 544 | 457 | 374 | 505 | 520 | 765 | 567 | 523 | 0.451002 | 0.370946 | 0.477988 | -0.08384 | -0.36576 |
| YMR045C | 14 | 17 | 26 | 50 | 13 | 15 | 1253 | 1493 | 3574 | 4335 | 1018 | 1079 | 0.011154 | 0.009312 | 0.01316 | -0.23861 | -0.49895 |
| RPL2A | 924 | 1092 | 394 | 844 | 1357 | 906 | 5394 | 5358 | 5890 | 7737 | 8290 | 6754 | 0.157776 | 0.080528 | 0.129471 | 0.285243 | -0.68507 |
| RPL2B | 6472 | 5942 | 4684 | 7374 | 7938 | 8872 | 3186 | 2880 | 2739 | 3951 | 4998 | 3900 | 0.671831 | 0.641069 | 0.65414 | 0.038497 | -0.02912 |
| RIM1 | 462 | 482 | 439 | 727 | 522 | 518 | 3931 | 3459 | 3411 | 4763 | 3230 | 3254 | 0.113736 | 0.123224 | 0.138227 | -0.28135 | -0.16575 |
| IST1 | 61 | 39 | 33 | 97 | 66 | 84 | 194 | 273 | 459 | 569 | 237 | 388 | 0.182108 | 0.106359 | 0.197894 | -0.11993 | -0.89578 |
| RPL39 | 2012 | 2755 | 1653 | 2310 | 2128 | 1885 | 4961 | 5006 | 5275 | 7402 | 6875 | 5900 | 0.321761 | 0.238224 | 0.239249 | 0.427476 | -0.0062 |
| AIM11 | 126 | 110 | 106 | 123 | 110 | 135 | 449 | 391 | 708 | 754 | 394 | 303 | 0.219346 | 0.135236 | 0.263237 | -0.26315 | -0.96088 |
| YBR255C-A | 215 | 146 | 255 | 277 | 269 | 147 | 317 | 237 | 364 | 442 | 312 | 202 | 0.392668 | 0.398606 | 0.442099 | -0.17106 | -0.14941 |
| RPL30 | 6540 | 7192 | 5352 | 8270 | 10648 | 9792 | 18610 | 17113 | 11145 | 14212 | 19069 | 14409 | 0.277973 | 0.346136 | 0.381462 | -0.4566 | -0.1402 |
| OAZ1 | 6 | 3 | 7 | 1 | 0 | 5 | 346 | 559 | 292 | 520 | 363 | 350 | 0.011192 | 0.012665 | 0.007042 | 0.668329 | 0.846781 |
| UBC4 | 487 | 516 | 781 | 631 | 631 | 836 | 6677 | 6154 | 6756 | 6786 | 6256 | 5168 | 0.07267 | 0.094348 | 0.115431 | -0.6676 | -0.29096 |
| EFB1 | 3566 | 4324 | 2957 | 4451 | 3976 | 3729 | 39560 | 42117 | 39259 | 53137 | 53145 | 47410 | 0.087898 | 0.073667 | 0.071263 | 0.302676 | 0.047879 |
| SEC27 | 575 | 808 | 511 | 676 | 595 | 565 | 3794 | 4260 | 4128 | 6007 | 4233 | 3812 | 0.14552 | 0.105653 | 0.126162 | 0.205948 | -0.25594 |
| RPL33B | 1088 | 1187 | 804 | 1357 | 1434 | 1140 | 17533 | 16158 | 13490 | 17122 | 24484 | 21241 | 0.063432 | 0.064841 | 0.053132 | 0.255617 | 0.287321 |
| PTC7 | 382 | 493 | 387 | 598 | 607 | 424 | 725 | 635 | 760 | 902 | 600 | 448 | 0.391067 | 0.368034 | 0.494569 | -0.33876 | -0.42633 |
| RPL33A | 4693 | 4812 | 3193 | 4110 | 5007 | 4094 | 37134 | 35248 | 29923 | 35680 | 43726 | 35567 | 0.11616 | 0.099855 | 0.102984 | 0.173691 | -0.04451 |
