## Supplementary material for "Revisiting the window of opportunity for co-transcriptional splicing efficiency and fidelity"

| | premRNA_fast_A | premRNA_fast_B | premRNA_slow_A | premRNA_slow_B | premRNA_wt_A | premRNA_wt_B | mRNA_fast_A | mRNA_fast_B | mRNA_slow_A | mRNA_slow_B | mRNA_wt_A | mRNA_wt_B | premRNA ratio_fast | premRNA ratio_slow | premRNA ratio_wt | premRNA ratio_fast/wt(log2) | premRNA ratio_slow/wt(log2) |
|---|---|---|---|---|---|---|---|---|---|---|---|---|---|---|---|---|---|
| RPL39 | 2012 | 2755 | 1653 | 2310 | 2128 | 1885 | 4961 | 5006 | 5275 | 7402 | 6875 | 5900 | 0.321761 | 0.238224 | 0.239249 | 0.427476 | -0.0062 |
| RPL40A | 11578 | 11571 | 9516 | 12981 | 15202 | 11219 | 12834 | 13138 | 13110 | 17679 | 19349 | 15927 | 0.471283 | 0.421982 | 0.426635 | 0.143589 | -0.01582 |
| RPL40B | 1335 | 1314 | 810 | 1473 | 1725 | 1690 | 16250 | 16269 | 15322 | 19020 | 22130 | 19643 | 0.075324 | 0.061044 | 0.075766 | -0.00844 | -0.31169 |
| RPL42A | 11151 | 11837 | 12080 | 16110 | 16205 | 16179 | 5470 | 4413 | 4676 | 5711 | 6610 | 5455 | 0.699665 | 0.729608 | 0.729064 | -0.05938 | 0.001075 |
| RPL42B | 2253 | 2601 | 1548 | 2454 | 2372 | 2549 | 14555 | 13001 | 8923 | 11540 | 16536 | 15128 | 0.150376 | 0.161599 | 0.134824 | 0.157499 | 0.261339 |
| RPL43A | 8885 | 8716 | 5671 | 7026 | 8053 | 7478 | 14611 | 14106 | 10330 | 13222 | 15620 | 12328 | 0.380031 | 0.350706 | 0.358869 | 0.082657 | -0.0332 |
| **RPL43B** | 3053 | 3135 | 3584 | 4954 | 4634 | 3965 | 2577 | 2367 | 1921 | 2675 | 2807 | 2542 | 0.556033 | 0.650204 | 0.616055 | -0.14789 | 0.077835 |
| RPL6A | 1869 | 1736 | 941 | 1284 | 1294 | 1219 | 10228 | 9716 | 8809 | 12512 | 13318 | 11724 | 0.153045 | 0.094792 | 0.09137 | 0.744169 | 0.053043 |
| RPL6B | 1880 | 2029 | 1415 | 1676 | 2143 | 1619 | 18820 | 19719 | 15785 | 20311 | 26280 | 21313 | 0.092059 | 0.079247 | 0.072998 | 0.334688 | 0.118495 |
| RPO26 | 388 | 518 | 656 | 800 | 774 | 761 | 1785 | 1731 | 2349 | 2777 | 2945 | 2348 | 0.20444 | 0.220977 | 0.226447 | -0.1475 | -0.03528 |
| RPP1B | 3459 | 3348 | 2721 | 4154 | 4685 | 5164 | 61825 | 60268 | 45962 | 65371 | 62118 | 59455 | 0.052806 | 0.05782 | 0.075023 | -0.50663 | -0.37576 |
| RPS0A | 2248 | 2230 | 1165 | 1730 | 2037 | 1714 | 20993 | 18887 | 17594 | 23039 | 23527 | 20927 | 0.101164 | 0.065974 | 0.077693 | 0.38084 | -0.23587 |
| RPS0B | 3436 | 3944 | 2107 | 2795 | 4120 | 2971 | 28071 | 25287 | 17741 | 25125 | 24649 | 24511 | 0.12199 | 0.103132 | 0.125658 | -0.04274 | -0.28501 |
| RPS10A | 2958 | 3108 | 2017 | 3432 | 3742 | 3886 | 39842 | 37670 | 35960 | 43875 | 45573 | 40250 | 0.072665 | 0.062829 | 0.081963 | -0.17371 | -0.38353 |
| RPS10B | 2235 | 2413 | 1848 | 2647 | 2051 | 2076 | 30938 | 27425 | 24684 | 28147 | 24466 | 21717 | 0.074122 | 0.077805 | 0.0823 | -0.15098 | -0.08102 |
| RPS11A | 2284 | 2349 | 1378 | 2189 | 2451 | 2033 | 44844 | 40867 | 41816 | 49592 | 60786 | 47667 | 0.051409 | 0.037088 | 0.039832 | 0.368095 | -0.10297 |
| RPS11B | 2947 | 3426 | 1945 | 2912 | 3469 | 2996 | 23216 | 21589 | 19059 | 23283 | 29458 | 25085 | 0.124799 | 0.101884 | 0.106023 | 0.235231 | -0.05745 |
| RPS13 | 32757 | 33391 | 13861 | 20174 | 25585 | 21921 | 12713 | 12960 | 10449 | 14866 | 17348 | 14839 | 0.720402 | 0.572959 | 0.596128 | 0.273179 | -0.05719 |
| RPS14A | 5304 | 6061 | 1692 | 2339 | 2619 | 2663 | 31736 | 26432 | 22857 | 27777 | 29542 | 27726 | 0.164865 | 0.073295 | 0.084532 | 0.963708 | -0.20579 |
| **RPS14B** | 3614 | 3997 | 2598 | 3947 | 6078 | 5713 | 3418 | 2511 | 1744 | 1988 | 4471 | 3473 | 0.564052 | 0.63169 | 0.599047 | -0.08684 | 0.076548 |
| RPS16A | 2909 | 3290 | 2137 | 3259 | 2995 | 3108 | 9436 | 10175 | 9242 | 13327 | 13092 | 12440 | 0.23999 | 0.192147 | 0.193036 | 0.314101 | -0.00666 |
| RPS16B | 2875 | 2873 | 1517 | 2319 | 2851 | 2554 | 24775 | 24828 | 18867 | 24463 | 29972 | 26523 | 0.103846 | 0.080505 | 0.087348 | 0.249609 | -0.1177 |
| RPS17A | 8545 | 7274 | 6069 | 7438 | 9703 | 8882 | 17861 | 17134 | 15351 | 20995 | 26661 | 21157 | 0.310809 | 0.272465 | 0.281256 | 0.144144 | -0.04581 |
| RPS17B | 2307 | 2659 | 1159 | 1809 | 2513 | 2020 | 4141 | 4492 | 4621 | 5765 | 5737 | 4407 | 0.364811 | 0.219681 | 0.309453 | 0.23743 | -0.49431 |
| RPS18A | 2935 | 2986 | 2103 | 3215 | 3425 | 3251 | 29420 | 29306 | 23611 | 33687 | 30949 | 33262 | 0.091591 | 0.084453 | 0.094338 | -0.04264 | -0.15968 |
| RPS18B | 1113 | 1204 | 971 | 1151 | 1045 | 1252 | 23529 | 24150 | 21514 | 31723 | 30896 | 32434 | 0.046327 | 0.039098 | 0.034942 | 0.40691 | 0.162161 |
| **RPS19A** | 2397 | 2393 | 1629 | 2372 | 2452 | 2225 | 16794 | 16373 | 13905 | 17881 | 15652 | 15929 | 0.12621 | 0.110993 | 0.129001 | -0.03156 | -0.21692 |
| RPS19B | 2845 | 2911 | 1683 | 2603 | 2719 | 2406 | 25124 | 25245 | 22186 | 28165 | 27950 | 25344 | 0.102554 | 0.077555 | 0.08768 | 0.226072 | -0.17701 |
| RPS21A | 2681 | 2194 | 1338 | 2088 | 2601 | 2657 | 31967 | 32994 | 19657 | 27747 | 29812 | 27827 | 0.069865 | 0.066857 | 0.083703 | -0.26072 | -0.3242 |
| **RPS21B** | 886 | 935 | 744 | 1058 | 1233 | 1217 | 28754 | 29480 | 20819 | 30790 | 31625 | 28061 | 0.030317 | 0.033862 | 0.039546 | -0.38342 | -0.22387 |
| RPS22B | 4582 | 4571 | 2017 | 3779 | 5328 | 5360 | 14320 | 13836 | 8298 | 9625 | 11326 | 9231 | 0.245369 | 0.238736 | 0.343636 | -0.48593 | -0.52547 |
| RPS23A | 2079 | 2254 | 1471 | 2387 | 2630 | 2454 | 23914 | 21271 | 18446 | 26058 | 23788 | 21631 | 0.087898 | 0.078886 | 0.100721 | -0.19647 | -0.35252 |
| RPS23B | 2643 | 2903 | 1728 | 2786 | 2817 | 2721 | 47937 | 48544 | 34209 | 46954 | 46921 | 45832 | 0.05434 | 0.052048 | 0.056339 | -0.05212 | -0.11431 |
| RPS24A | 6335 | 7118 | 3099 | 4826 | 4982 | 4635 | 10687 | 10767 | 8193 | 10148 | 10496 | 9874 | 0.385076 | 0.298367 | 0.320667 | 0.26407 | -0.10399 |
| RPS24B | 1499 | 1589 | 885 | 1749 | 2196 | 1859 | 4255 | 3174 | 2362 | 3619 | 3516 | 3342 | 0.297064 | 0.299189 | 0.370943 | -0.32042 | -0.31014 |
| RPS27A | 3377 | 4145 | 2529 | 3496 | 4226 | 3003 | 1673 | 1383 | 1033 | 1231 | 1554 | 1447 | 0.709266 | 0.724788 | 0.702987 | 0.012829 | 0.044061 |
| RPS27B | 1964 | 2719 | 1359 | 2193 | 2391 | 1937 | 16178 | 15957 | 15716 | 23960 | 24736 | 24605 | 0.126923 | 0.081721 | 0.08056 | 0.655815 | 0.020653 |
| RPS30A | 3479 | 4608 | 1919 | 3777 | 2680 | 2778 | 23433 | 22750 | 17665 | 21855 | 21698 | 15351 | 0.148853 | 0.122672 | 0.131585 | 0.177894 | -0.1012 |
| RPS30B | 1416 | 1744 | 1087 | 2059 | 1452 | 1471 | 7751 | 7140 | 5288 | 8186 | 8001 | 7222 | 0.175388 | 0.185743 | 0.161409 | 0.119822 | 0.202584 |
| RPS4A | 1222 | 1514 | 825 | 1253 | 1516 | 1494 | 24674 | 22558 | 21900 | 26360 | 29382 | 23970 | 0.055042 | 0.04084 | 0.053868 | 0.0311 | -0.39943 |
| RPS4B | 1363 | 1384 | 561 | 1058 | 1308 | 996 | 13148 | 13605 | 8297 | 13005 | 12833 | 11929 | 0.093132 | 0.069283 | 0.084778 | 0.135572 | -0.2912 |

| | premRNA_fast_A | premRNA_fast_B | premRNA_slow_A | premRNA_slow_B | premRNA_wt_A | premRNA_wt_B | mRNA_fast_A | mRNA_fast_B | mRNA_slow_A | mRNA_slow_B | mRNA_wt_A | mRNA_wt_B | premRNA ratio_fast | premRNA ratio_slow | premRNA ratio_wt | premRNA ratio_fast/wt(log2) | premRNA ratio_slow/wt(log2) |
|---|---|---|---|---|---|---|---|---|---|---|---|---|---|---|---|---|---|
| RPS6A | 1627 | 1788 | 1264 | 2138 | 1999 | 1861 | 6254 | 6308 | 6391 | 9254 | 8148 | 8710 | 0.213648 | 0.176398 | 0.186526 | 0.195859 | -0.08054 |
| RPS6B | 4319 | 4714 | 2009 | 3067 | 3773 | 3055 | 13989 | 16458 | 11840 | 18193 | 17903 | 16307 | 0.22928 | 0.144663 | 0.165923 | 0.466594 | -0.19782 |
| RPS7A | 3545 | 3785 | 2635 | 3514 | 3450 | 3321 | 77457 | 76820 | 48960 | 68065 | 74992 | 65096 | 0.045361 | 0.050082 | 0.046261 | -0.02835 | 0.114486 |
| RPS7B | 2022 | 1952 | 1568 | 2366 | 3173 | 2610 | 27095 | 23675 | 21254 | 22776 | 30048 | 24798 | 0.072807 | 0.081406 | 0.09537 | -0.38946 | -0.2284 |
| RPS9A | 4568 | 4763 | 3262 | 4822 | 5716 | 6217 | 212 | 219 | 839 | 975 | 996 | 849 | 0.955845 | 0.813613 | 0.865728 | 0.142863 | -0.08957 |
| RPS9B | 8394 | 9030 | 4237 | 5585 | 9919 | 8361 | 25405 | 24513 | 18985 | 22399 | 27688 | 24705 | 0.258779 | 0.191017 | 0.258306 | 0.002637 | -0.43538 |
