## Supplementary material for "Revisiting the window of opportunity for co-transcriptional splicing efficiency and fidelity"

**Supplemental Information**

**Table S2**

Transcripts with non-consensus 5'SS whose splicing efficiency was measurable in RNA-seq data from [26], indicating those with improved (green) or reduced (red) splicing efficiency in the fast mutant compared to WT. The list of genes with non-consensus 5'SS sequences was obtained using the search facility of the Ares Lab Yeast Intron Database: http://intron.ucsc.edu/yeast4.3/, and checked in the Saccharomyces Genome Database: https://www.yeastgenome.org/

### Genes with non-canonical 5’ splice sites and their splicing efficiency

Of 24 RPs (including YML6*, a mitochondrial RP) with non-canonical 5’SS, 19 splice worse (red) and 5 splice better (green) with fast RNAPII
Of 12 non-RPs with the same non-canonical 5’SS, 4 splice worse and 8 splice better with fast RNAPII
The non-RPs are colour coded in the 1st column. The RPs are colour coded in the 2nd column

**5’SS GTACGT**: 19 introns: (including **14 RPs**, plus YML6*, a mitochondrial RP)

| ORF | Aliases | position | 5’ SS | Branch Point | 3’ SS |
|---|---|---|---|---|---|
| YBR189W | RPS9B | chrII:604509-604932 | GTACGT | **TGACTAAC**AAGA | TTTTAG |
| YDL012C | | chrIV:431467-431391 | GTACGT | **GTACTAAC**GAGC | ATTCAG |
| YDR447C | RPS17B | chrIV:1355545-1355241 | GTACGT | **TTACTAAC**TTAA | TTATAG |
| YDR471W | RPL27B | chrIV:1401795-1402189 | GTACGT | **TTACTAAC**AAAT | TCGTAG |
| YFL034C-A | RPL22B | chrVI:64915-64604 | GTACGT | **TTACTAAC**ATAA | TAACAG |
| YGL226C-A | OST5 | chrVII:73132-72993 | GTACGT | **TTACTAAC**AAGT | CTTAAG |
| YHR010W | RPL27A | chrVIII:126546-127117 | GTACGT | **TTACTAAC**TATT | ACACAG |
| YIL133C | RPL16A | chrIX:99380-99100 | GTACGT | **ATACTAAC**AAAT | AACCAG |
| YJL136C | RPS21B | chrX:157244-156794 | GTACGT | **GTACTAAC**ATTG | TACTAG |
| YJL191W | RPS14B | chrX:73791-74209 | GTACGT | **TTACTAAC**AACT | GTATAG |
| YLR287C-A | RPS30A | chrXII:713150-712730 | GTACGT | **ATACTAAC**ATGG | ATTTAG |
| YLR306W | UBC12 | chrXII:744148-744292 | GTACGT | **ATACTAAC**ATAA | AAATAG |
| YML025C | YML6* | chrXIII:225333-225244 | GTACGT | **ATACTAAC**ATTA | CAATAG |
| YMR033W | ARP9 | chrXIII:337812-337908 | GTACGT | **ATACTAAC**AACA | TAACAG |
| YNL069C | RPL16B | chrXIV:494968-494529 | GTACGT | **TTACTAAC**TTTA | TTATAG |
| YNL302C | RPS19B | chrXIV:62918-62377 | GTACGT | **TTACTAAC**AAAA | AATTAG |
| YOL121C | RPS19A | chrXV:92825-92445 | GTACGT | **TTACTAAC**AATA | CTACAG |
| YOR182C | RPS30B | chrXV:678785-678384 | GTACGT | **TTACTAAC**ATTA | CCTTAG |
| YPL081W | RPS9A | chrXVI:404951-405462 | GTACGT | **TCACTAAC**AATG | AAACAG |

**5’SS GTATGC**: 8 introns, including **2 RPs**

| ORF | Aliases | position | 5’ SS | Branch Point | 3’ SS |
|---|---|---|---|---|---|
| YBL059W | | chrII:110874-110953 | GTATCG | **TTACTAAC**ATAA | ACATAG |
| YCL002C | | chrIII:111628-111562 | GTATGC | TT**GACTAAC**TAA | ACATAG |
| YDR305C | HNT2 | chrIV:1073396-1073317 | GTATGC | **TTACTAAC**TATA | TTACAG |
| YEL003W | GIM4 PFD2 | chrV:148189-148287 | GTATGC | CT**TACTAAC**AAT | GTATAG |
| YIL148W | RPL40A | chrIX:68710-69154 | GTATGC | **TTACTAAC**TTGT | TAACAG |
| YJR094W-A | RPL43B | chrX:608301-608586 | GTATGC | **TTACTAAC**AAAT | TTGCAG |
| YLL050C | COF1 | chrXII:40395-40226 | GTATGC | **TTACTAAC**TAAT | GTGTAG |
| YNL050C | YNL050C | chrXIV:534960-534879 | GTATGC | **TTACTAAC**TTTT | TTGCAG |

**5’SS GTATGA**: 6 introns in our data (**5 RPs**)

| ORF | Aliases | position | 5’ SS | Branch Point | 3’ SS |
|---|---|---|---|---|---|
| YBR048W | RPS11B | chrII:332870-333391 | GTATGA | **TTACTAAC**ATGC | ATACAG |
| YDR025W | RPS11A | chrIV:491554-491903 | GTATGA | **TTACTAAC**CTTT | TTTTAG |
| YIL004C | BET1 | chrIX:348489-348368 | GTATGA | **TTACTAAC**TATA | ATATAG |
| YLR061W | RPL22A | chrXII:263200-263599 | GTATGA | **TTACTAAC**ATTA | ATATAG |
| YPL090C | RPS6A | chrXVI:378384-378000 | GTATGA | A**TACTAAC**AAAT | AAATAG |
| YPR043W | RPL43A | chrXVI:654162-654575 | GTATGA | **TTACTAAC**ATAT | ATACAG |

**5’SS GTCNNN**: **1 RP**

| ORF | Aliases | position | 5’ SS | Branch Point | 3’ SS |
|---|---|---|---|---|---|
| YGL030W | RPL30 | chrVII:439088-439328 | GTCAGT | C**TACTAAC**AAGT | CAACAG |

**5’SS GTGNNN**: 2 introns, including **1 RPG**

| ORF | Aliases | position | 5’ SS | Branch Point | 3’ SS |
|---|---|---|---|---|---|
| YML034W | SRC1 | chrXIII:211439-211575 | GTGAGT | **TTACTAAC**ATCT | CTCTAG |
| YMR242C | RPL20A | chrXIII:754214-753747 | GTGAGT | CA**TACTAAC**ATT | TATCAG |

### The following non-canonical 5’SS are not found in RPs

**5’SS GTAGTA**: 1 intron – **no RPs**

| ORF | Aliases | position | 5’ SS | Branch Point | 3’ SS |
|---|---|---|---|---|---|
| YCL012C | | chrIII:101695-101638 | GTAGTA | A**TACTAAC**AGCT | TAATAG |

**5’SS GTAANN**: 11 introns – **no RPs.** **N.B. GTAAGT stabilises interaction with U1 snRNA**
Of 11 for which we have data (but low counts, so not all reliable), 3 splice worse and 8 splice better with fast RNAPII. Some also have non-consensus BP sequences (red)

| ORF | Aliases | position | 5’ SS | Branch Point | 3’SS |
|---|---|---|---|---|---|
| YAL030W | SNC1 | chrI:87382-87505 | GTAAGT | A**TACTAAC**TTTC | TTTTAG |
| YBL059C-A | | chrII:110500-110425 | GTAAGT | **TTACTAAC**ATTG | TTTTAG |
| YDL064W | UBC9 | chrIV:337519-337639 | GTAAGT | A**TACTAAC**AAAT | TAACAG |
| YFR045W | | chrVI:242004-242086 | GTAAGT | TT**TATTAAC**GCT | TTGCAG |
| YHR001W-A | QCR10 | chrVIII:107826-107899 | GTAAGT | A**TACTAAC**ATTT | ATACAG |
| YIL156W-A | YIL156W-B | chrIX:47693-47765 | GTAAGT | AA**TACTAAC**AAG | GGTTAG |
| YKR095W-A | PCC1 | chrXI:625896-625981 | GTAAGT | **TTACTAAC**TATT | CATTAG |
| YLR211C | | chrXII:564508-564459 | GTAAGT | T**GACTAAC**ATGA | AATTAG |
| YLR445W | | chrXII:1024567-1024659 | GTAAGT | TT**TACTAAT**AAT | TAACAG |
| YNL012W | SPO1 | chrXIV:609785-609879 | GTAAGT | A**AACTAAC**CGAA | TATTAG |
| YNL044W | YIP3 | chrXIV:545286-545375 | GTAAGT | A**TACTAAC**GCGT | CTATAG |
